# Down syndrome with Alzheimer’s disease brains have increased iron and associated lipid peroxidation consistent with ferroptosis

**DOI:** 10.1101/2025.02.05.636731

**Authors:** Max A. Thorwald, Jose A. Godoy-Lugo, Elizabeth Kerstiens, Gilberto Garcia, Minhoo Kim, Sarah J. Shemtov, Justine Silva, Salma Durra, Peggy A. O’Day, Wendy J. Mack, Annie Hiniker, Marc Vermulst, Bérénice A. Benayoun, Ryo Higuchi-Sanabria, Henry Jay Forman, Elizabeth Head, Caleb E. Finch

## Abstract

**INTRODUCTION:** Cerebral microbleeds (MB) are associated with sporadic Alzheimer’s Disease (AD) and Down Syndrome with AD (DSAD). Higher MB iron may cause iron mediated lipid peroxidation. We hypothesize that amyloid deposition is linked to MB iron and that amyloid precursor protein (APP) triplication increases iron load and lipid peroxidation.

**METHODS:** Prefrontal cortex and cerebellum of cognitively normal (CTL), AD and DSAD ApoE3,3 carriers were examined for proteins that mediated iron metabolism, antioxidant response, and amyloid processing in lipid rafts.

**RESULTS:** Iron was 2-fold higher in DSAD than CTL and AD. Iron storage proteins and lipid peroxidation were increased in prefrontal cortex, but not in the cerebellum. The glutathione synthesis protein GCLM was decreased by 50% in both AD and DSAD. Activity of lipid raft GPx4, responsible for membrane repair, was decreased by at least 30% in AD and DSAD.

**DISCUSSION:** DSAD shows greater lipid peroxidation than AD consistent with greater MBs and iron load.

**Graphical Abstract:** 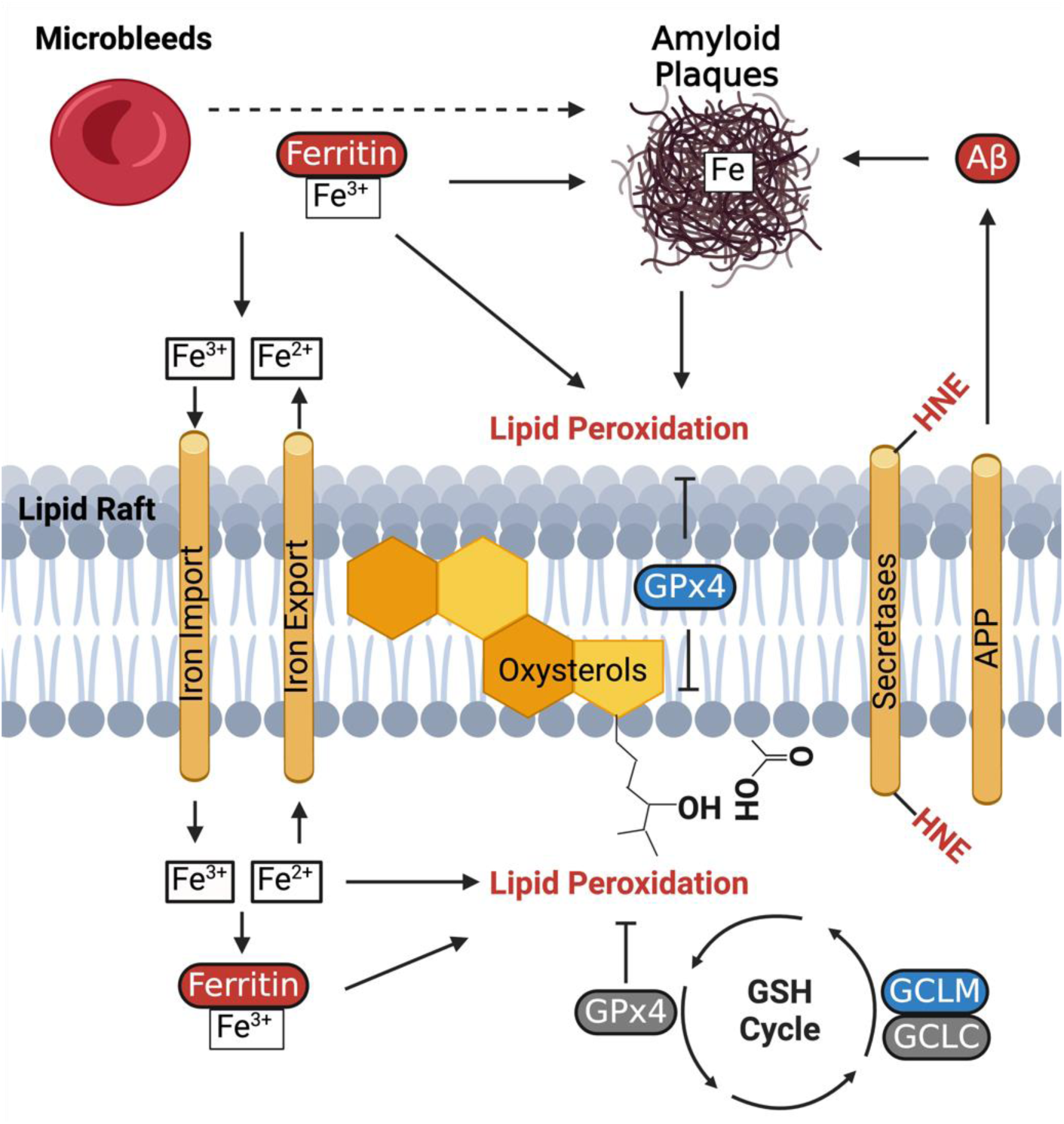

Cerebral microbleeds result in increased brain iron and lipid peroxidation in DSAD consistent with ferroptosis as reported for Alzheimer’s disease. Aβ; beta-amyloid peptides, APP; amyloid precursor protein, GCLC; glutathione cysteine ligase catalytic subunit, GCLM; glutathione cysteine modifier subunit, GPx4; glutathione peroxidase 4, HNE; 4-hydroxynonenal.

**RESEARCH IN CONTEXT:**

1. **Systematic Review:** DS is associated with increased microbleeds and brain iron that may be mediated by increased APP from Trisomy 21. To assess potential links between amyloid and iron levels, we examined sporadic and DS with AD brains for amyloid processing and antioxidant enzyme defense in lipid rafts. We further compared DSAD with rare variants of DS: partial and mosaic T21.
2. **Interpretation:** DSAD brains showed greater oxidation of lipid rafts where APP is processed than sporadic AD. Corresponding decreases in lipid raft antioxidant enzymes, despite increased total levels of these antioxidant enzymes, present a new mechanism for aberrant amyloid processing during AD.
3. **Future Directions:** Iron chelation therapies in combination with amyloid monoclonals may benefit DSAD.

## INTRODUCTION

Down Syndrome (DS) afflicts one in seven hundred births each year because of the triplication of chromosome 21[1]. Trisomy 21 (T21) confers increased gene dosage of the amyloid precursor protein (APP), and other proteins (BACE2, S100β, DYRK1A, RCAN1) involved in Alzheimer’s disease (AD) pathology. APP is processed in lipid rafts by the secretase enzymes [2–4]. APP is cleaved by the non-amyloidogenic α-secretases (ADAM family) or pro-amyloidogenic β-secretases (BACE1/2) yielding soluble amyloid beta peptides (Aβ) from processing by the γ-secretase complex. Soluble Aβ42, and to a lesser extent Aβ40, are aggregated into cytotoxic oligomers implicated in neuronal death. Soluble and oligomeric Aβ progressively form higher order protofibrils, fibrils, and ultimately extracellular senile plaques. The increased production of Aβ peptides and increased β-secretase activity with age are considered the primary drivers of AD [5]. Sporadic AD is the most common form of AD, which increasingly occurs after age 60 in all apolipoprotein E (ApoE) genotypes [6]. However, in DS, this timeframe is shifted earlier, with the onset of AD and cognitive impairment by age 40-50 [7], half of DS individuals are AD-positive by age 60 [8]. The increase in APP gene dosage and earlier onset of AD in DS individuals supports the role of Aβ as a driver of AD.

Cerebral microbleeds (MBs) may be the main source of increased brain iron in all forms of AD: sporadic, autosomal dominant, or familial AD (FAD), and DSAD. MBs also occur in pediatric cases of DS, particularly in the frontal lobes [9,10]. ApoEFAD mice, containing human FAD transgenes and human ApoE, developed ‘Naked’ MBs, absent of amyloid plaque localization at 2 months of age, independent of sex and ApoE genotype. Subsequently, these MBs increase with age and are found to colocalize with both amyloid plaques and activated microglia by immunohistochemistry [11]. In most human imaging studies, MBs are harder to characterize due to the limited availability of high-resolution MRI. Sporadic AD may have hundreds of MBs [12], which may be further increased in DSAD and FAD [9,13]. Cerebral amyloid angiopathy (CAA) is strongly implicated in MBs FAD and DSAD [9,13]. While there is evidence of retrograde amyloid transport from parenchyma to cerebral arteries, the converse has not been indicated [14]. For these reasons, we do not further consider CAA. Collectively these observations implicate MBs and the resulting deposition of iron in the formation of amyloid plaques.

Microglia degrade extravasated blood from MB in the brain parenchyma [15]. Additionally, erythrocytes lyse readily by oxidation, releasing hemoglobin [16]. The breakdown of extracellular hemoglobin by microglia sometimes results in cell death due to the high iron load leaving an iron-laden hemosiderin deposit comprised of heme, iron-loaded ferritin, and other iron rich proteins at the site of MBs [17–19]. The ferrous iron (Fe^2+^) from heme can oxidize lipid membranes in surrounding cells (**Graphical Abstract**). Furthermore, heme can be imported into cells by heme carrier protein 1 (HCP1), particularly in neuronal and glial cells, which possess higher levels of HCP1. Intracellular heme is degraded by heme oxygenase (HMOX) to liberate Fe^2+^ which is converted to non-reactive ferric iron (Fe^3+^) for storage transport by transferrin or for storage by ferritin. Increases in cellular iron is tightly regulated to inhibit oxidative damage to proteins, DNA, and lipids. Ferrous iron can undergo Fenton chemistry, causing lipid peroxidation in AD [20], by reacting with hydrogen peroxide (H_2_O_2_), yielding a potent hydroxyl radical (**^·^**OH). Hydroxyl radicals react instantaneously to oxidize polyunsaturated lipids in their immediate environment [21]. These oxidized lipids decompose to a variety of products including carbonyls such as α,β-unsaturated hydroxyalkenal 4-hydroxynonenal (HNE), which form adducts with amino acids cysteine, histidine, and lysine in proteins [22]. These reactions from **^·^**OH with membrane lipids occur rapidly due to the neighboring iron-rich hemosiderin deposits from MBs and extra-cellular amyloid plaques. Amyloid plaques are also rich in iron and copper [23,24]. Therefore, MB iron has a large role in oxidative damage of cellular membranes and APP processing.

APP processing occurs in lipid rafts (LR), which are small hydrophobic microdomains present in cell plasma membranes of many cell types and enriched in cholesterol, sphingomyelin, and phosphatidylcholine. These tightly packed domains facilitate signal transduction and if disrupted cause cell death [25,26]. Due to their high content of lipids with double bonds, such as arachidonic acid, lipid rafts are easily oxidized [27]. Reduced capability of antioxidant enzyme systems or increased iron-mediated lipid peroxidation results in iron-mediated cell death, often referred to as ferroptosis [28]. The most abundant non-enzymatic antioxidant is glutathione (GSH) used by several critical enzymes to safeguard against ferroptosis. These GSH-dependent enzymes remove oxidants including hydrogen peroxide, lipid hydroperoxides, and electrophilic oxidation products including HNE preventing pathogenic modifications of membrane proteins or lipids within the lipid raft. Recently we confirmed the presence of ferroptotic markers in sporadic AD human prefrontal cortex with concomitant increases in lipid raft oxidation, reduced GSH-reliant antioxidant enzymes, and increased iron storage [29]. Intriguingly, BACE1 protein levels and enzyme activity are increased by lipid peroxidation [30,31]. Due to the increase of MBs with DS [9,10,32], we hypothesize that ferroptosis-related changes will be increased in DSAD due to the higher APP gene dosage. Brains from cognitively normal, sporadic AD, and DSAD are compared in the AD impacted prefrontal cortex and the ‘AD resistant’ cerebellum for changes in iron signaling, lipid peroxidation, and amyloid processing as related to ferroptosis and AD. These studies address the gaps in data on DS lipid rafts and APP processing and antioxidant defense.

## METHODS

### Human Samples

Postmortem prefrontal cortex and cerebellum were provided by Alzheimer’s Disease Centers at the University of Southern California and the University of California Irvine (UCI MIND). Tissues were matched for ApoE3,3 to remove the contributions of ApoE allele differences on AD neuropathology. Samples were genotyped for ApoE alleles by PCR for SNP variants rs429358 and rs7412. An equal number of both sexes were included (n=4/sex/group). All human subjects provided informed consent. Human tissue use was approved through IRB protocol #UP-20-00014-EXEMPT. Details for individual brains are in **Supplemental Table 1**.

### Tissue Washing

Blood in brain tissue was minimized by washing [29]. Frozen tissues were thawed on ice and minced with surgical scissors. Samples were washed twice with PBS by vortexing and then centrifuged at 800g/30 sec/4°C for further analysis.

### Lipid Raft

40 mg of human brain tissue was used for isolation of lipid rafts from prefrontal cortex and cerebellum using a commercial kit (Invent Biotechnologies, Plymouth, MN). LRs were previously validated in comparison to traditional ultracentrifuge methods for brain tissue and cells [29,33].

### Biochemical Assays

Total protein was quantified by 660nm assay (Thermo Fisher Scientific, Waltham, MA). Tissue heme was quantified by Quantichrom assay (BioAssay Systems, Hayward, CA). Total glutathione peroxidase activity was calculated by activity assay (Cayman Chemical, Ann Arbor, MI). For phospholipid hydroperoxidase activity, oxidized phosphatidyl choline (PCOOH) was generated as previously reported and used in place of the provided cumene hydroperoxide [29]. The same batch of PCOOH was utilized for all assays. Cholesterol assays were performed on lipid raft fractions according to the manufacturer’s protocol (Cell Biolabs, San Diego, CA). Secretase activity assays were performed as previously described [34,35]. Briefly, lipid raft lysates were adjusted to pH4.5 for which maximum activity has been reported [36,37]. Förster resonance energy transfer substrates containing the a- (10uM) or β-cleavage (20uM) sites for APP protein (Sigma, St. Louis, MO) were mixed with LR lysates. The kinetic assays were carried out at 37°C for 60 cycles of 60 seconds. LR lysates were validated to have the highest amount of a- and β-secretase activity compared to the non-raft membrane (**Supplemental Figure 1**). All optical densities were measured using a SpectraMax M2 spectrophotometer equipped with a temperature regulator (Molecular Devices, San Jose, CA).

### Western Blot

whole cell lysates derived from radioimmunoprecipitation buffer (20ug) were boiled at 75°C under denatured conditions and resolved on 4-20% gradient gels. Proteins were electroblotted using a Criterion blotter (Bio-Rad Laboratories, Hercules, CA) and transferred onto 0.45um polyvinyl difluoride membranes. Membranes were stained using Revert 700 fluorescent protein stain and imaged before blocking with LI-COR Intercept blocking buffer (LI-COR Biosciences, Lincoln, NE), followed by primary antibodies. Membranes incubated with IRDye 800CW and/or 700CW secondary antibodies and visualized by Odyssey (LI-COR Biosciences). Western blot data was quantified with ImageJ and normalized by total protein per lane and/or loading control protein.

### ICP-MS

50 mg of brain tissue was cut using a ceramic scalpel and placed in metal-free test tubes. Tissues were washed in PBS twice and homogenized with sterilized disposable plastic pestles in Chelex 100 (Sigma, St. Louis, MO) treated purified water (18.2 MΩ; Millipore, Burlington, MA). Homogenates were desiccated by vacuum centrifuge at 95°C for 90 min. Desiccated pellets were dissolved in trace metal-free 70% HNO_3_ overnight. 30% H_2_O_2_ was then added the remaining solution was boiled off. The remaining pellet was resuspended in 2% HNO_3_ and analyzed by Agilent 7500ce ICP-MS in hydrogen mode with a practical detection limit of 10ppb and a relative standard deviation (RSD) of replicate measures between 0.2 to 5%. Total iron concentration was normalized to wet-weight tissue.

### RNA sequencing

20 mg of brain tissue was homogenized in TRIzol reagent using a BeadBug Benchtop Homogenizer. Brain tissue was suspended in 1 mL of TRIzol and homogenized for 6 rounds of 10 s homogenization and 60-second holds between each round. 300 µL of chloroform was added to the sample and aqueous separation of RNA was performed using centrifugation in a heavy gel phase-lock tube (VWR, 10847-802). The aqueous phase was applied to a standard column-based RNA purification kit (Quantabio, Extracta Plus, Cat# 95214-050) following the manufacturer’s protocol. Library preparation (mRNA library, poly A enrichment) and RNA sequencing (NovoSeq PE150. 6G raw data per sample) was performed by Novogene. Reads were trimmed with trim_galore-0.6.5-1 and mapped to WBcel235 with STAR-2.7.3a [38]. Mapped reads were counted to genes using featureCounts (Subread-2.0.3) [39]. Artifacts were removed with RUVSeq-1.32.0 [40] using R-4.3.2 and differential expression analysis was performed with DESeq2-1.34.0 [41]. Raw RNA-seq data is available through Annotare: E-MTAB-14179.

### Statistics

ANCOVA was performed for all data using SPSS (Ver. 29.0.20.0; Chicago, IL) to adjust for sex differences among study groups. Pairwise comparisons were made using Bonferroni (p<0.05). Levene’s test was used to test for homogeneity of variances. Non-normal data underwent log transformation. Correlation matrixes used Spearman correlations for non-normal data.

## RESULTS

The brain tissues were defined for AD based on Braak staging based on location and abundance of neurofibrillary tangles (NFT). All cognitively normal had Braak scores 0-1, 3-5 for AD, and 6 for DSAD. The average age of death for cognitively normal was 89 years and 85 years for AD. DSAD had over a two-decade decrease in average age of death at 58 years, independent of sex (**S.Table 1**).

### APP Triplication is associated with increased iron and lipid peroxidation in the prefrontal cortex

T21 increases the gene dosage of APP and other AD-relevant genes. APP protein levels were 2.5-fold higher in DSAD prefrontal cortex than in cognitively normal and in AD brains (**Fig. 1A**), consistent with APP triplication. Total iron levels were 2-fold higher in DSAD than in both cognitively normal and AD (**Fig. 1B**). Lipid peroxidation product HNE, which is formed from iron-catalyzed lipid peroxidation, was increased in DSAD relative to CTL 65% and AD 50% (**Fig. 1C**).

**Figure 1:**
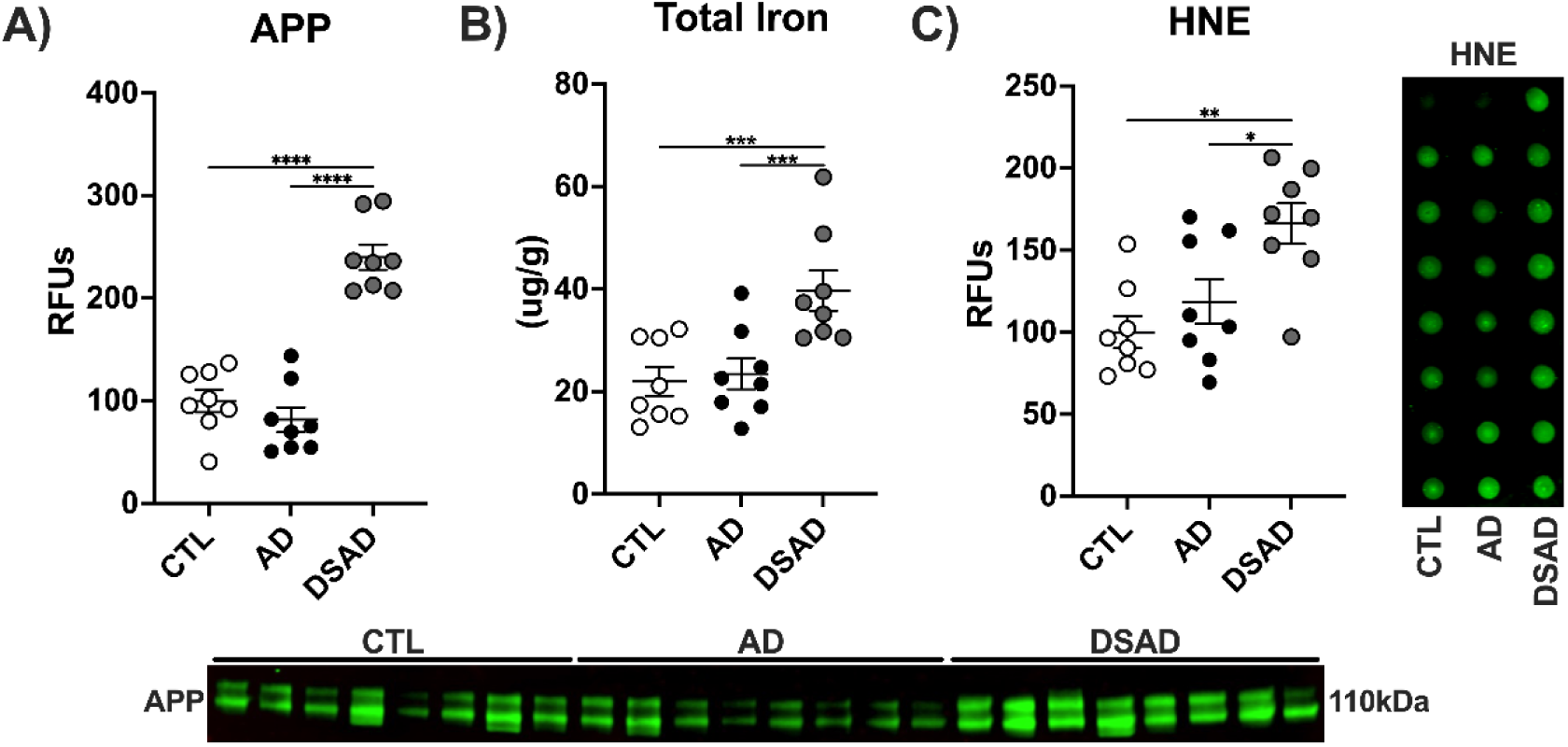
APP gene dosage is associated with iron and lipid peroxidation in DSAD. Western blots for **A)** APP, **B)** Total iron by ICP-MS. **C)** HNE by dot blot. Data are shown as relative fluorescent units (RFUs) for western or dot blots. Statistics by ANCOVA adjusted for sex with Bonferroni’s posthoc test. *p<0.05, **p<0.01, ***p<0.001, ****p<0.0001.

**Extended Figure 1:**
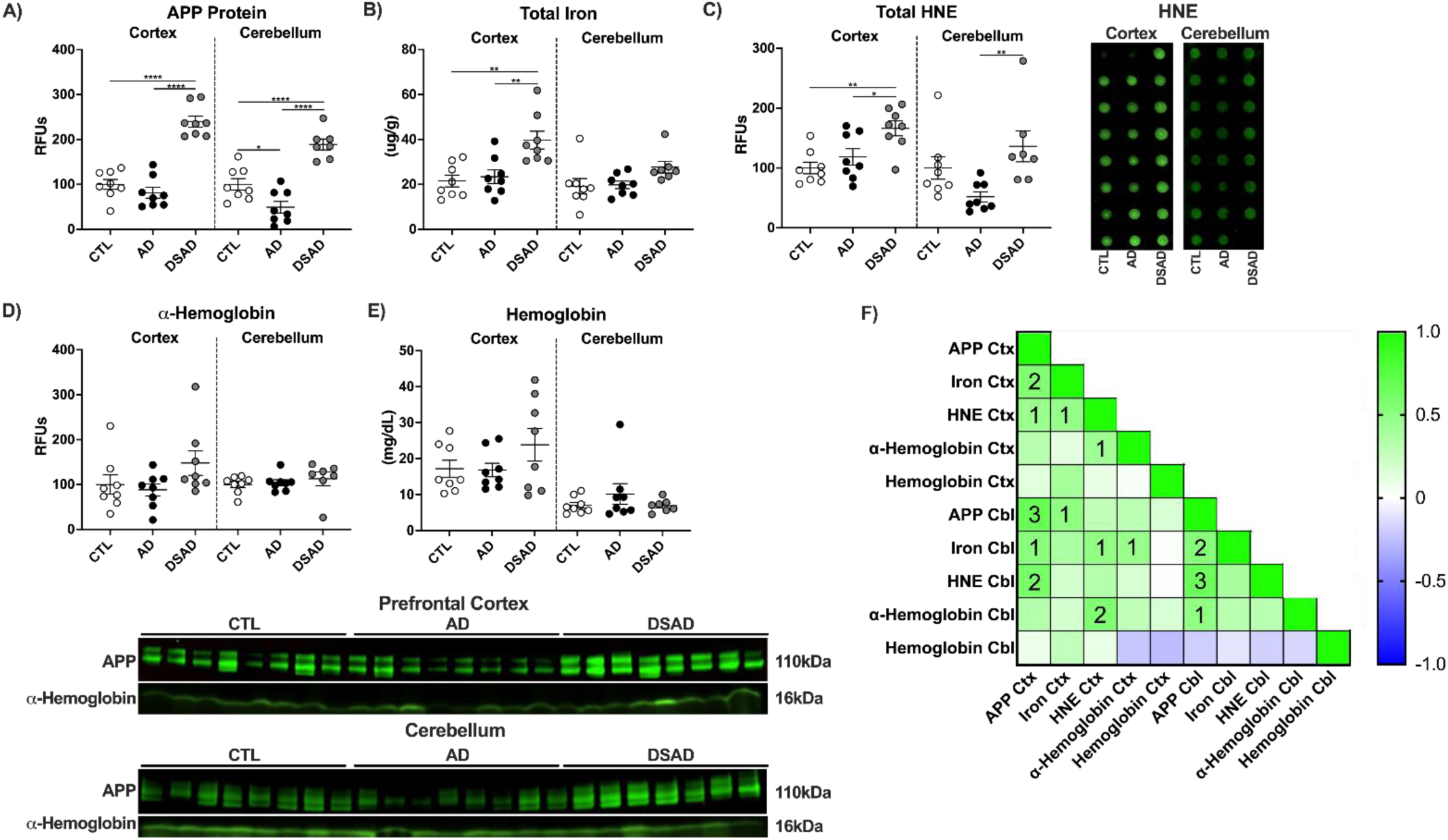
APP gene dosage is associated with iron and lipid peroxidation in DSAD brains. **A)** APP, **B)** Total Iron, **C)** HNE, **D)** ⍺-hemoglobin, and **E)** tissue hemoglobin. **F)** Correlation matrix for prefrontal cortex (Ctx) and cerebellum (Cbl). Data are shown as RFUs from western or dot blots. Statistics by ANCOVA adjusted for sex with Bonferroni’s posthoc test. *p<0.05, **p<0.01, ***p<0.001, ****p<0.0001. Correlation matrix analyzed by Spearman correlation. 1p<0.05, 2p<0.01, 3p<0.001, 4p<0.0001.

Prefrontal cortex and cerebellum were compared (**Ext. Fig. 1A**). Cerebellum APP decreased by 50% in AD but was almost 2-fold higher with DSAD compared to cognitively normal (**Ext. Fig. 1A**). Total iron did not differ in cerebellum (**Ext. Fig. 1B**). HNE increased 1.5-fold in DSAD above AD (**Ext. Fig. 1C**). No differences were observed for ⍺-hemoglobin or tissue hemoglobin levels in either brain region (**Ext. Fig. 1D,E**).

APP, total iron, and HNE were all positively correlated with each other in prefrontal cortex. These relationships were conserved in cerebellum except for HNE and total iron (**Ext. Fig. 1F**).

### How is iron metabolism altered in relation to brain MBs

Having shown that DSAD had increased total iron in prefrontal cortex but not the cerebellum, we examined iron signaling and metabolism (**Fig. 2A**). Transferrin (TF) transports the majority of ferric iron through blood to cells by internalization after binding to the Transferrin receptor (TfR). In the prefrontal cortex, TfR, but not TF, increased by over 80% in DSAD compared to cognitively normal and AD brains (**Fig. 2B,C**). Ferrous iron is imported by divalent metal transporter 1 (DMT1) or exported by ferroportin 1 (FPN). DMT1 and FPN were not different in prefrontal cortex (**Fig. 2D,E**). Iron also enters the cell as heme derived from erythrocyte breakdown through heme carrier protein 1 (HCP1). HCP1 decreased by 35% in prefrontal cortex of AD and DSAD (**Fig. 2F**). Heme is degraded by hemeoxygenase (HMOX) to ferrous iron for storage as ferritin. HMOX1 levels increased 3-fold in DSAD compared to both cognitively normal and AD (**Fig. 2G**) implying an increase in erythrocyte breakdown from MBs. Neuron-specific HMOX2 was decreased by 45% in AD below cognitively normal (**Fig. 2H**). Excess imported iron is stored by the ferritin complex comprised of ferritin light (FTL) and heavy (FTH1) chains. Iron storage protein FTL increased 3.5-fold in DSAD compared to cognitively normal and 2-fold compared to AD (**Fig. 2I**). FTH1 levels were 3-fold higher in DSAD prefrontal cortex compared to CTL and AD (**Fig. 2J**). Iron regulatory proteins (IRP) regulate the mRNA transcripts of these iron metabolism genes. The IRP proteins did not differ between DSAD, AD, and CTL (**Fig. 2K,L**).

**Figure 2:**
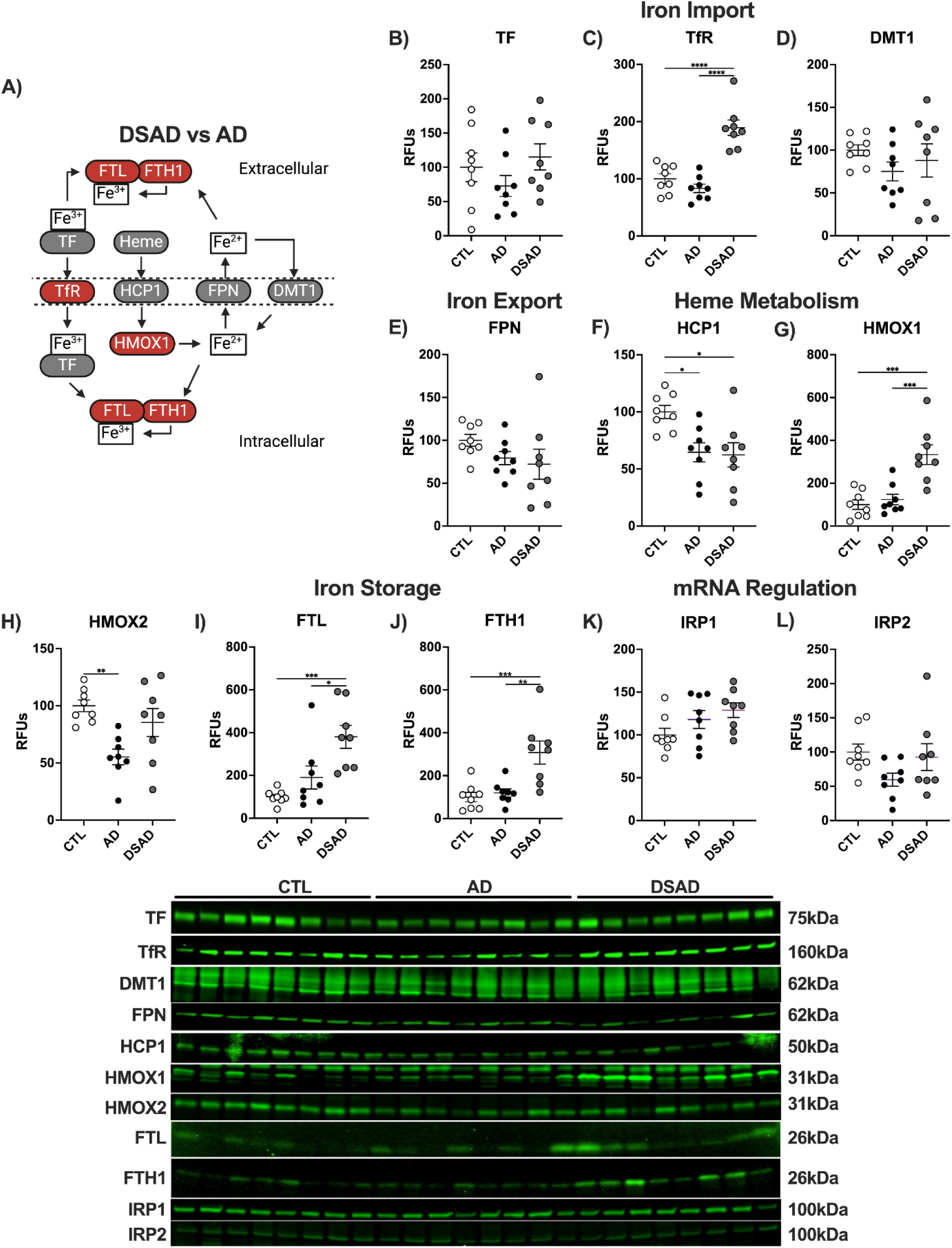
Iron signaling and storage proteins increase with AD. **A)** Schema for iron signaling changes (red, increase; grey, no change; blue, decrease) with DSAD versus AD. Western blots are shown as relative fluorescent units for human prefrontal cortex in RIPA lysates for **B)** TF, **C)** TfR, **D)** DMT1, **E)** FPN, **F)** HCP1, **G)** HMOX1, **H)** HMOX2, **I)** FTL, and **J)** FTH1, **K)** IRP1, and **L)** IRP2. Statistics by ANCOVA adjusted for sex with Bonferroni’s posthoc test. Correlation matrix by Spearman correlation. *p<0.05, **p<0.01, ***p<0.001, ****p<0.0001.

**Extended Figure 2:**
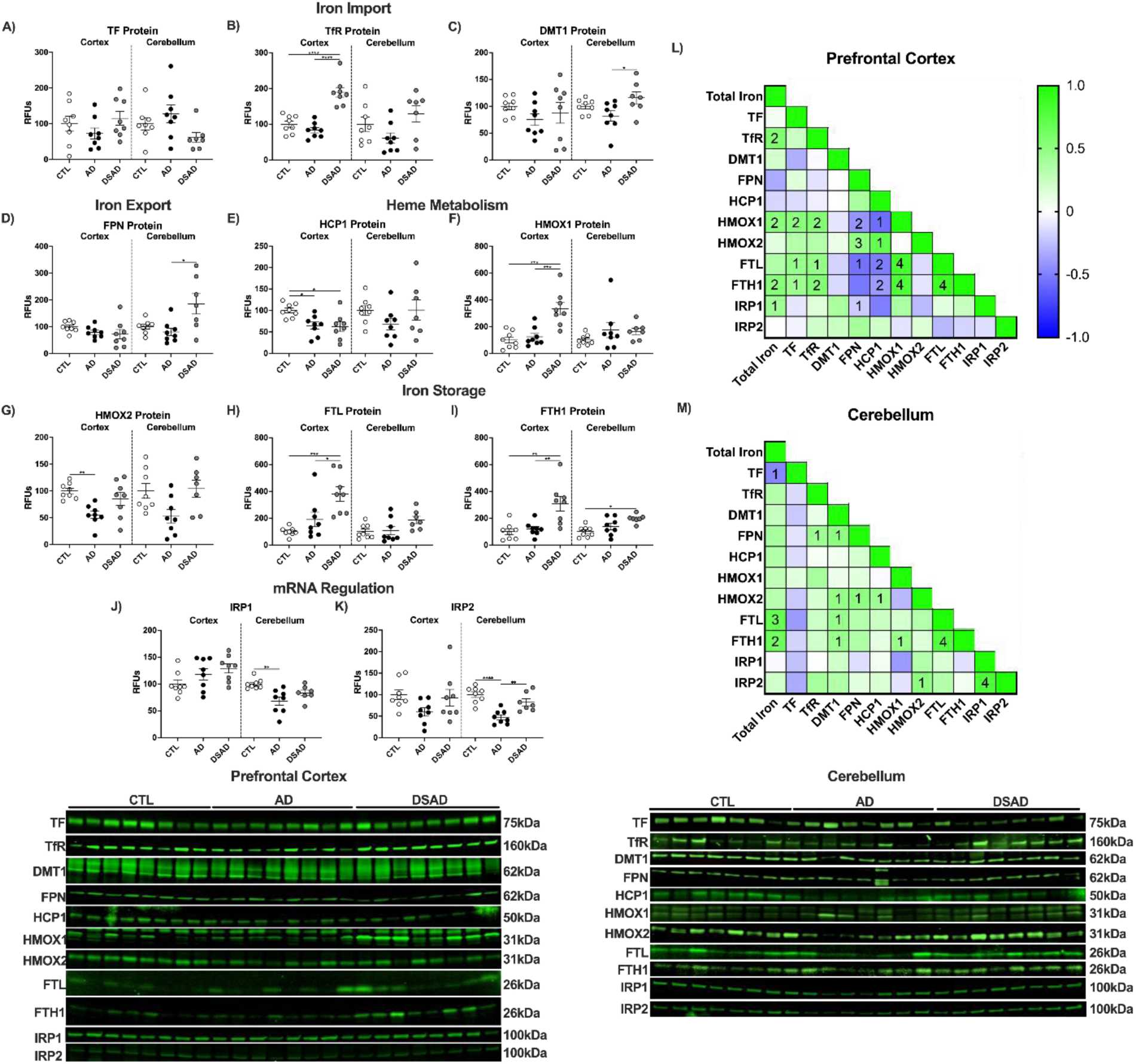
Iron signaling and storage proteins increase with AD. Western blots are shown as relative fluorescent units for human prefrontal cortex and cerebellum in RIPA lysates for **A)** TF, **B)** TfR, **C)** DMT1, **D)** FPN, **E)** HCP1, **F)** HMOX1, **G)** HMOX2, **H)** FTL, **I)** FTH1, **J)** IRP1, and **K)** IRP2. Correlation matrixes for iron signaling proteins for **L)** prefrontal cortex and **M)** cerebellum. Statistics by ANCOVA adjusted for sex with Bonferroni’s posthoc test. *p<0.05, **p<0.01, ***p<0.001, ****p<0.0001. Correlation matrix analyzed by Spearman correlation. 1p<0.05, 2p<0.01, 3p<0.001, 4p<0.0001.

In the cerebellum, TF and TfR were differ with DSAD (**Ext. Fig 2A,B**), whereas DMT1 increased 40% (**Ext. Fig. 2C**). FPN increased 2-fold in DSAD (**Ext. Fig. 2D**). HCP1 was unaltered in cerebellum unlike prefrontal cortex (**Ext. Fig. 2E**). HMOX1 did not between the groups (**Ext. Fig. 2F**). HMOX2 increased 65% in DSAD cerebellum above AD (**Ext. Fig. 2G**). FTL did not differ in cerebellum while FTH1 increased 2-fold in DSAD cerebellum compared to CTL (**Ext. Fig. 2H, I**). IRP1 decreased 30% with AD compared to cognitively normal but did not differ for DSAD (**Ext. Fig. 2J**). IRP2 also decreased 50% with AD but was increased with DSAD 75% compared to AD (**Ext. Fig. 2K**). In prefrontal cortex HMOX1 was positively correlated with total iron, TF, and TfR, and the ferritin proteins, but were negatively correlated with HCP1 and FPN, suggesting the increased iron may be MB derived (**Ext. Fig. 2L**). These relationships were absent in the cerebellum (**Ext. Fig. 2M**).

### Antioxidant enzymes associated with mitigating ferroptosis through lipid radical detoxification are increased in DSAD

To combat oxidative damage derived from iron, we examined antioxidant enzymes relevant to lipid peroxidation and those relevant to DS such as SOD1 which resides on chromosome 21. The differences observed between AD and DSAD are outlined in **Figure 3A**. The glutathione peroxidase (GPx) and peroxiredoxin (Prdx) enzyme families are responsible for peroxide detoxification. GPx1 levels and total GPx activity did not differ between groups (**Fig. 3B,C**). Unique in the GPx and Prdx families are GPx4 and Prdx6 which reduce oxidized phospholipids and oxysterols. Prefrontal cortex GPx4 increased 2.5-fold in AD and DSAD compared to cognitively normal while Prdx6 was unaltered (**Fig. 3D,E**). Activity for GPx4 and Prdx6 measured by the reduction of oxidized phosphatidylcholine (PCOOH) was also unaltered in the prefrontal cortex (**Fig. 3F**). Quinone oxidation can result in superoxide (O_2_^·-^) production and Fenton chemistry. Membrane quinones are reduced by ferroptosis suppressor protein 1 (FSP1) while NADPH quinone oxidoreductase 1 (NQO1) reduces cytoplasmic quinones. FSP1 was increased by 80% in DSAD compared to AD and CTL (**Fig. 3G**). NQO1 was not altered in the prefrontal cortex (**Fig. 3H**). Decomposition of oxidized lipids results in reactive carbonyls which conjugate to proteins altering their structure and function. 4-hydroxynonenal (HNE) is the most abundant carbonyl formed from this process. The glutathione S-transferase (GST) or aldehyde reductase (ALDH) families’ clear carbonyls. GSH-dependent GSTA4 decreased in AD but not DSAD compared to cognitively normal while GSH-independent ALDH2 increased in DSAD compared to AD and CTL by at least 50% (**Fig. 3I,J**). Superoxide dismutase 1 (SOD1) which resides on chromosome 21 and reduces O_2_^·-^ was 2-fold higher in DSAD compared to both AD and CTL brains (**Fig. 3K**). These genes are regulated by transcription factors BTB domain and CNC homology 1 (BACH1) and Nuclear factor erythroid 2-related factor 2 (Nrf2). Nrf2 is the master regulator of antioxidant enzymes and iron metabolism genes. BACH1 is a competitive inhibitor of Nrf2 through competitive binding of Nrf2 promoters. Increases in heme degrade BACH1 allow Nrf2 to bind to antioxidant enzyme and iron promoters. Despite triplication of the BACH1 gene in DSAD, BACH1 protein decreased by 30% in DSAD compared to cognitively normal brains indicative of microbleed iron (**Fig. 3L**). Nrf2 protein was decreased by 45% in DSAD compared to cognitively normal (**Fig. 3M**).

**Figure 3:**
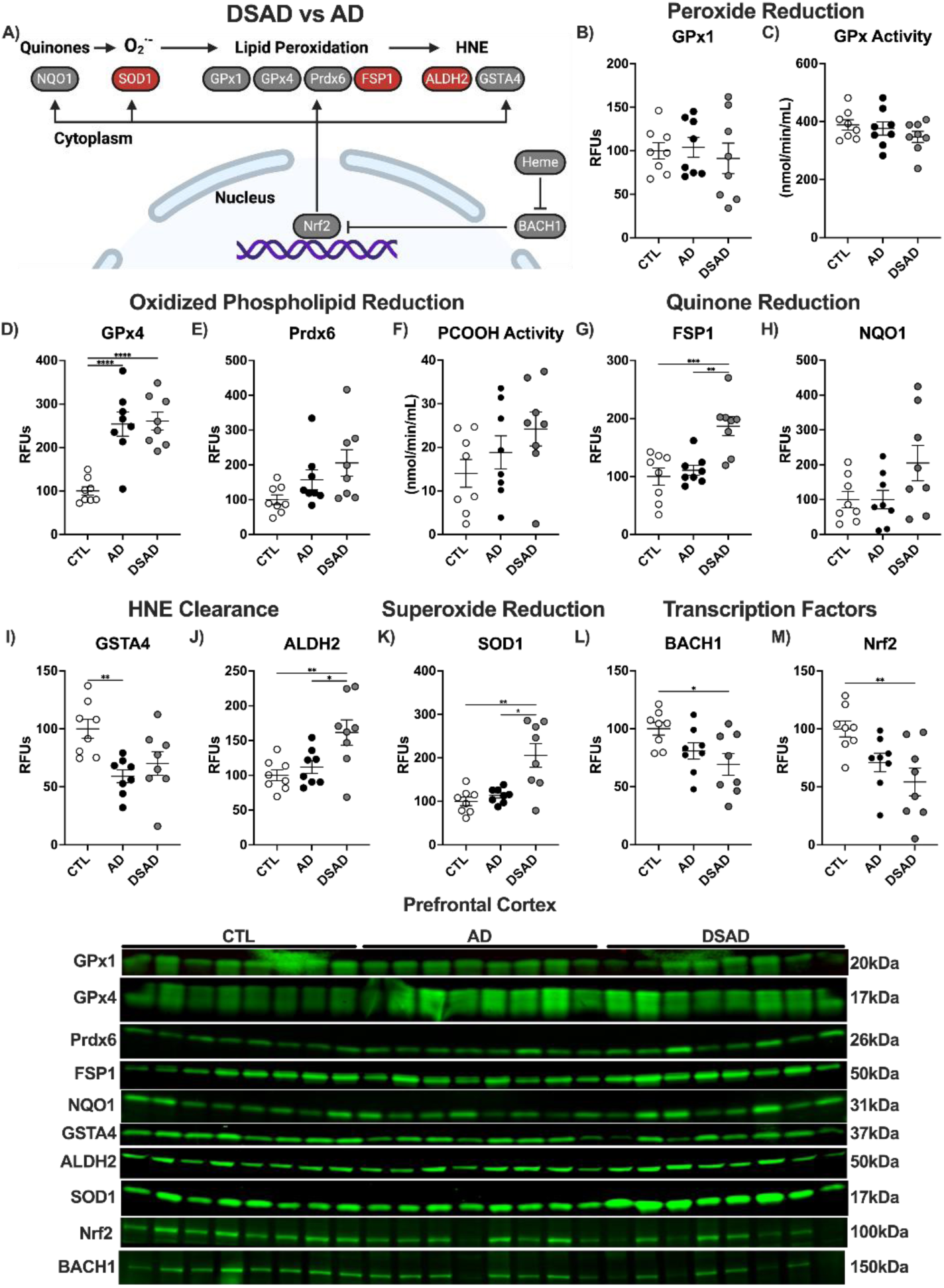
Antioxidant enzymes are selectively altered by DS with AD. **A)** Schematic representing antioxidant enzyme signaling changes (red increase, grey no change, blue decrease) with DSAD compared to AD. Western blots shown as relative fluorescent units or enzyme activities for human prefrontal cortex measured from RIPA lysates for **B)** GPx1, **C)** GPx Activity, **D)** GPx4, **E)** Prdx6, **F)** PCOOH activity, **G)** FSP1, **H)** NQO1, **I)** GSTA4, **J)** ALDH2, **K)** SOD1, **L)** BACH1, and **M)** Nrf2. Statistics by ANCOVA adjusted for sex with Bonferroni’s posthoc test. *p<0.05, **p<0.01, ***p<0.001, ****p<0.0001.

**Extended Figure 3:**
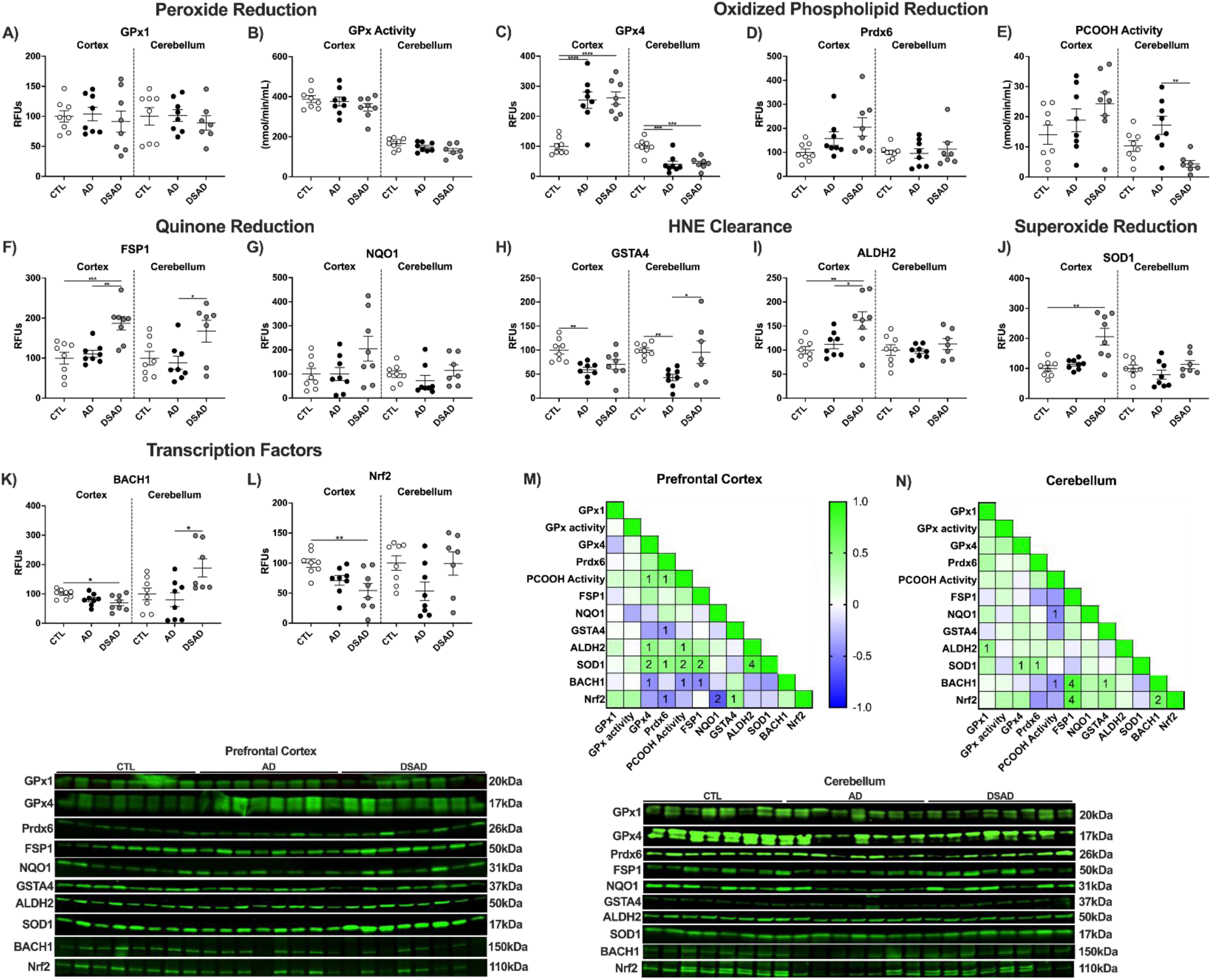
Antioxidant enzymes are altered by DS with AD. Western blots shown as relative fluorescent units or enzyme activities for human prefrontal cortex and cerebellum measured from RIPA lysates for **A)** GPx1, **B)** GPx activity, **C)** GPx4, **D)** Prdx6, **E)** PCOOH activity, **F)** FSP1, **G)** NQO1, **H)** GSTA4, **I)** ALDH2, **J)** SOD1, **K)** BACH1 and **L)** Nrf2. Correlation matrix of ferroptosis-related antioxidant enzymes for **M)** prefrontal cortex and **N)** cerebellum. Statistics by ANCOVA adjusted for sex with Bonferroni’s posthoc test. Correlation matrix analyzed by Spearman correlation. *p<0.05, **p<0.01, ***p<0.001, ****p<0.0001.

Cerebellum GPx1 and total GPx activity also did not differ between groups, paralleling the prefrontal cortex (**Ext. Fig. 3A,B**). In contrast with the prefrontal cortex, GPx4 protein decreased by at least 55% in the cerebellum of both AD and DSAD brains compared to cognitively normal (**Ext. Fig. 3C**). Prdx6 did not differ in the cerebellum (**Ext. Fig. 3D**). PCOOH activity decreased 75% in the cerebellum for DSAD brains compared to AD (**Ext. Fig. 3E**). FSP1 levels were elevated in DSAD brains by 90% compared to AD in cerebellum (**Ext. Fig. 3F**). No differences were observed in NQO1 (**Ext. Fig. 3G**). Cerebellum GSTA4 levels decreased with AD by 60% compared to cognitively normal, matching the prefrontal cortex while DSAD cerebellum had 125% more GSTA4 than AD (**Ext. Fig. 3H**). ALDH2 and SOD1 were unaltered in the cerebellum (**Ext. Fig. 3I,J**). BACH1 levels were 1.4-fold higher in DSAD compared to sporadic AD (**Ext. Fig. 3K**). Nrf2 levels did not differ in the cerebellum (**Ext. Fig. 3L**).

In prefrontal cortex, GPx4 and Prdx6 had positive correlations with PCOOH activity, consistent with their role in reducing phospholipid hydroperoxides (**Ext. Fig. 3K**). The cerebellum showed fewer correlations overall but showed a strong correlation between Nrf2 and FSP1 (**Ext. Fig. 3L**).

### GCLM, essential to GSH synthesis, is decreased in AD and DSAD

We next examined the GSH cycle, because GSH is essential to redox homeostasis and is required by many of these antioxidant enzymes (**Fig. 4A**). Cystine is the rate-limiting amino acid needed for GSH synthesis imported by the cystine/glutamate antiporter SLC7a11 (xCT). xCT works in conjunction with L-type amino acid transporter 1 (LAT1) which transports glutamine within somatic cells. Both cystine and glutamine are oxidized before ligation through the glutathione cysteine ligase (GCL) complex. Neither xCT nor LAT1 differed amongst cognitively normal, AD, or DSAD in the prefrontal cortex (**Fig. 4B,C**). GCL is comprised of a catalytic (GCLC) and modifier (GCLM) subunit which is the rate-limiting enzyme in GSH synthesis. GCLC did not differ while GCLM was decreased 45% in AD and 60% in DSAD compared to cognitively normal (**Fig. 4D,E**). The final step in GSH synthesis is the addition of glycine by glutathione synthetase (GSS) which also did not differ among the three groups (**Fig. 4F**). GSH is an electron donor for GPx, GST, Prdx, and other antioxidant enzyme families to reduce oxidants. The oxidation of GSH forms a disulfide bond between two tripeptides yielding GSSG. GSSG is converted back to GSH by glutathione reductase and NADPH. GSR, G6PD, and PGD levels were also unaltered in the prefrontal cortex with AD and DSAD compared to control (**Fig. 4G-I**).

**Figure 4:**
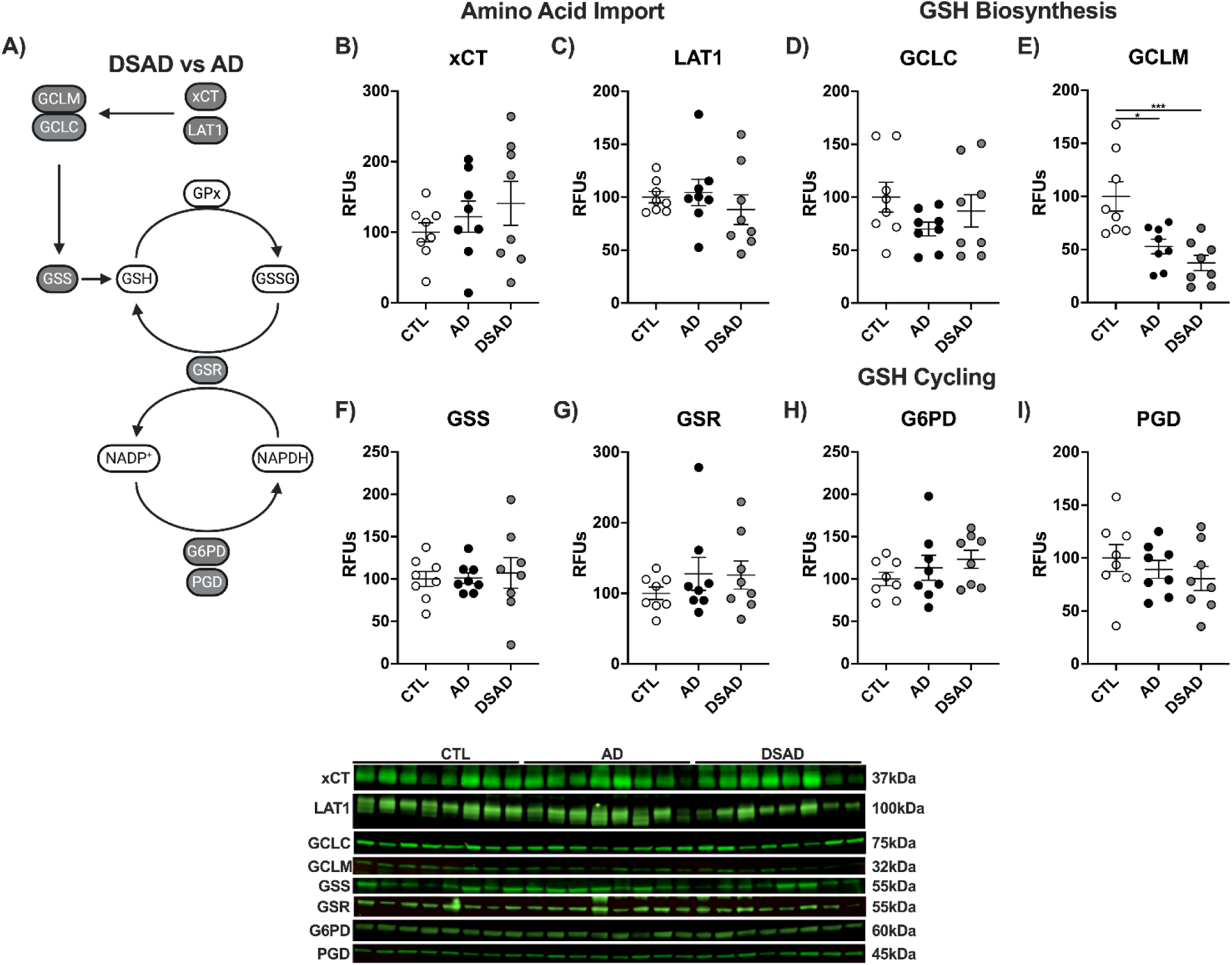
Decreased glutathione-producing enzyme GCLM with AD and DSAD. **A)** Proteins involved in the import of cystine, synthesis, and reduction of GSH. Western blots are shown as relative fluorescent units for **B)** xCT, **C)** LAT1, **D)** GCLC, **E)** GCLM, **F)** GSS, **G)** GSR, **H)** G6PD, and **I)** PGD in prefrontal cortex. Statistics by ANCOVA adjusted for sex with Bonferroni’s posthoc test. *p<0.05, ***p<0.001.

**Extended Figure 4:**
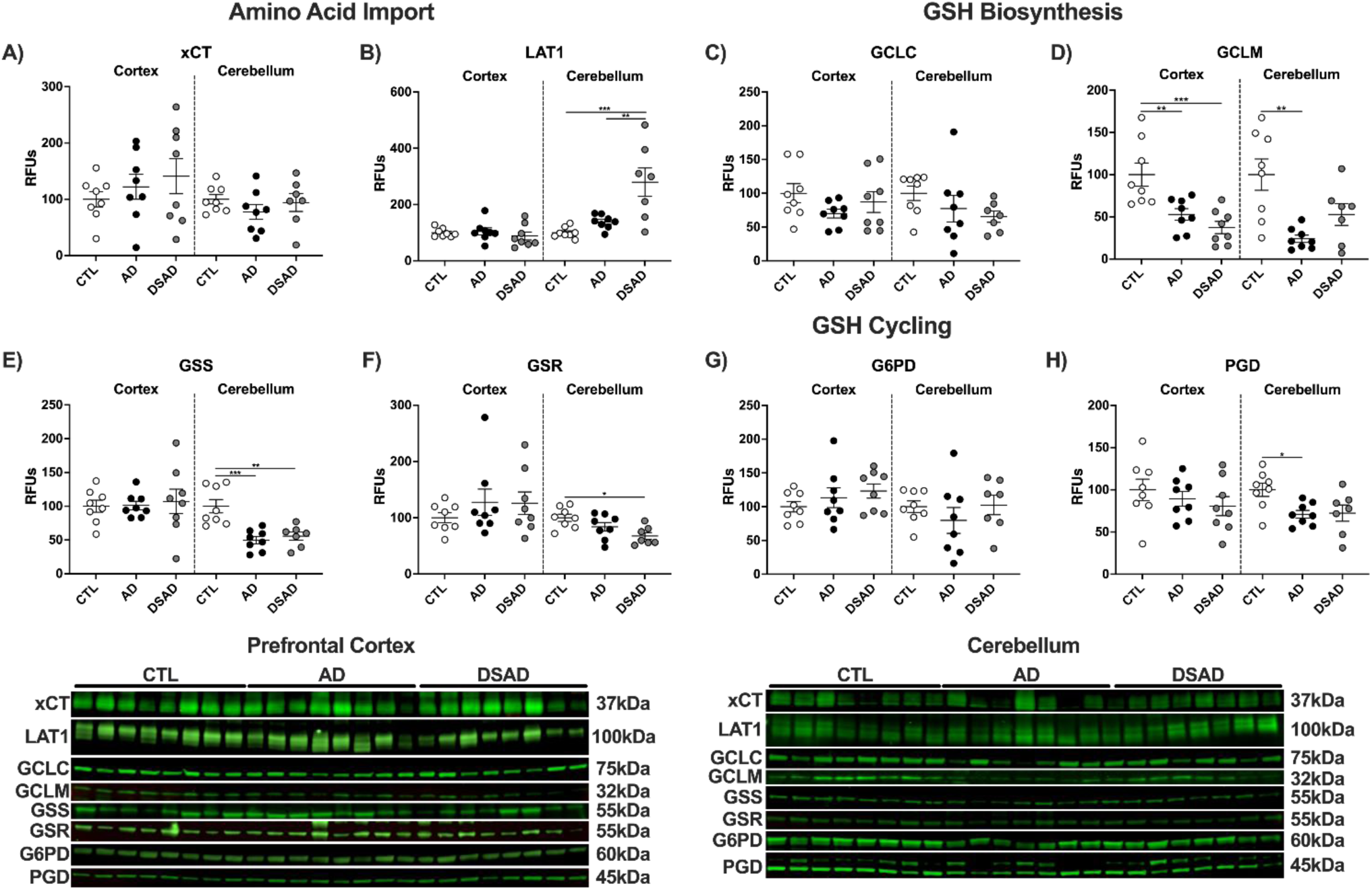
Loss of GCLM is consistent with AD and DSAD. **A)** Proteins involved in the import of cystine, synthesis, and reduction of GSH. Western blots are shown as relative fluorescent units for **B)** xCT, **C)** LAT1, **D)** GCLC, **E)** GCLM, **F)** GSS, **G)** GSR, **H)** G6PD, and **I)** PGD in prefrontal cortex. **J)** Representative images from Western blots. Statistics by ANCOVA adjusted for sex with Bonferroni’s posthoc test. *p<0.05, **p<0.01, ***p<0.001.

Cerebellum xCT did differ between the three groups (**Ext. Fig. 4A**), while LAT1 levels increased 3-fold in DSAD above CTL and AD (**Ext. Fig. 4B**). GCLC did not differ while GCLM was decreased by 75% in the AD cerebellum compared to CTL and DSAD (**Ext. Fig. 4C,D**). Cerebellum GSS levels were decreased by 50% in AD and 45% in DSAD compared to CTL (**Ext. Fig. 4E**). GSR protein was decreased by 30% in DSAD brains compared to CTL (**Fig. Ext. Fig. 4F**). G6PD was unaltered in the cerebellum while PGD was decreased with AD by 30% from CTL (**Ext. Fig. 4G,H**). Collectively these data suggest that GSH synthesis is impaired in both brain regions in AD and DSAD compared to CTL by low GCLM levels.

### Production of Aβ increases with DSAD

We next examined the enzymes that produce β-amyloid peptides Aβ40 and Aβ42 from APP in two phases. The initial phase analyzed whole cell lysates commonly done for pathological studies, followed by isolation of lipid rafts, the site of APP processing. APP is cleaved by the non-amyloidogenic (ADAM) or amyloidogenic (BACE) secretase enzymes to yield c-terminal fragments. These fragments are then processed by the γ-secretase complex which includes the catalytic subunit presenilin 1 (PSEN1) to produce Aβ peptides (**Fig. 5A**). ADAM10 increased 2.5-fold in DSAD prefrontal cortex compared to cognitively normal (**Fig. 5B**) while the amyloidogenic BACE1 did not differ between groups (**Fig. 5C**). PSEN1 decreased with AD by 45% but did not differ in DSAD relative to CTL (**Fig. 5D**).

**Figure 5:**
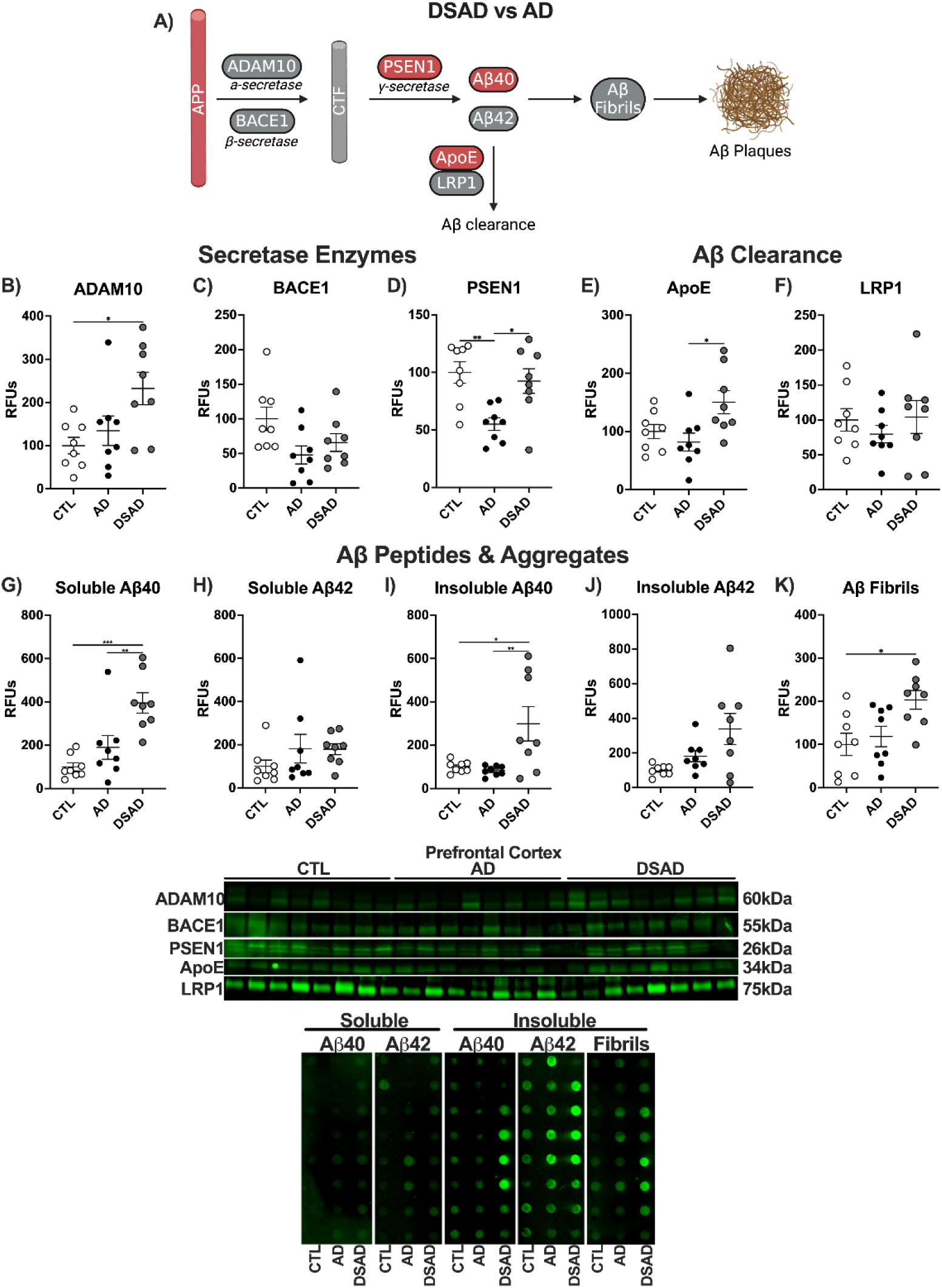
APP processing, clearance, and peptides in DSAD and AD. **A)** Differences in amyloid processing in DSAD compared to AD. Western blots are shown as relative fluorescent units for human prefrontal cortex measured from RIPA lysates for **B)** ADAM10, **C)** BACE1, **D)** PSEN1, **E)** ApoE, and **F)** LRP1. Dot blots for soluble **G)** Aβ40, **H)** Aβ42, insoluble **I)** Aβ40, **J)** Aβ42, and **K)** fibrillar amyloid. Statistics by ANCOVA adjusted for sex with Bonferroni’s posthoc test. *p<0.05, **p<0.01, ***p<0.001.

**Extended Figure 5:**
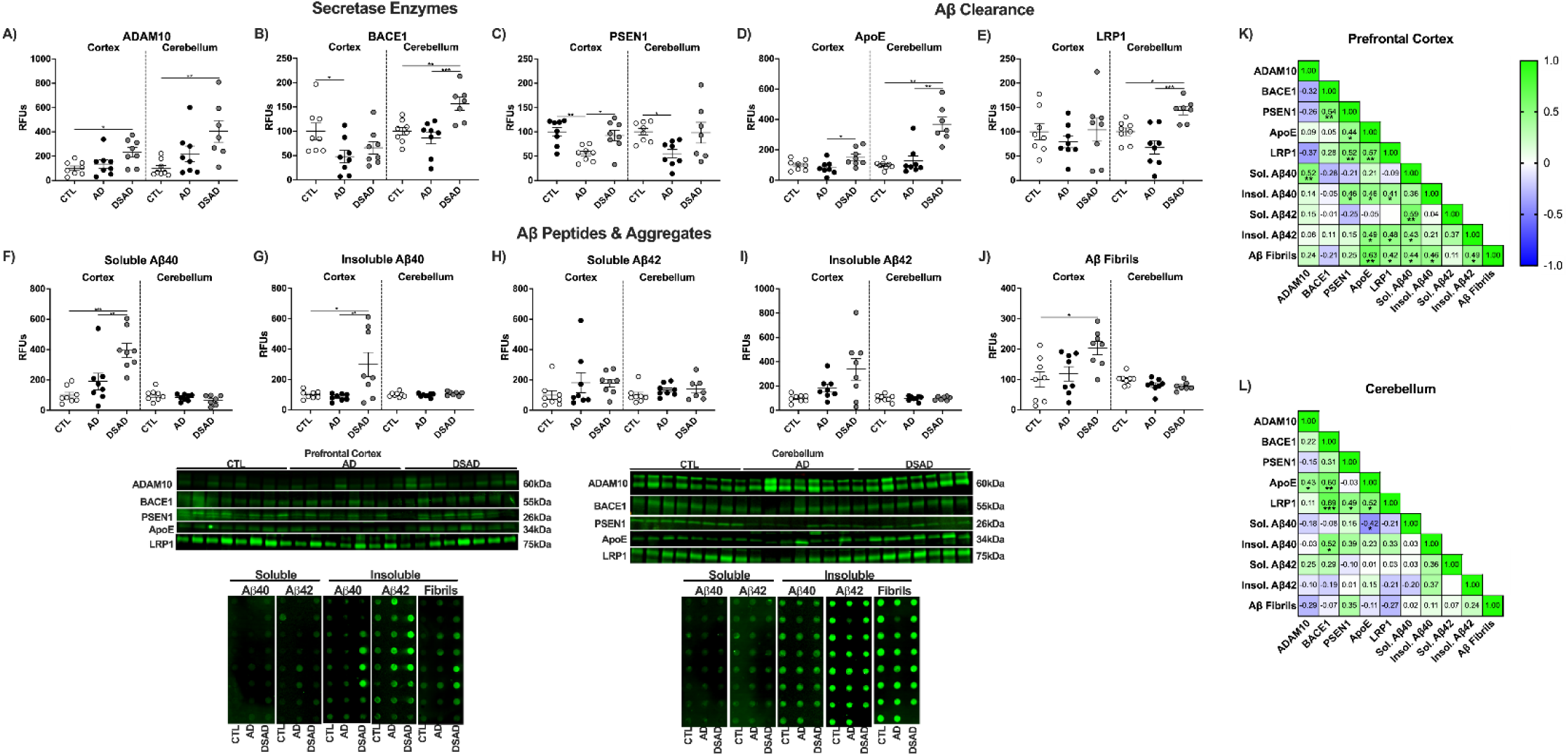
Amyloid processing, clearance, and peptides in DSAD and AD. Western blots are shown as relative fluorescent units for human prefrontal cortex and cerebellum measured from RIPA lysates for **A)** ADAM10, **B)** BACE1, **C)** PSEN1, **D)** ApoE, and **E)** LRP1. Dot blots for soluble **F)** Aβ40, **G)** Aβ42, insoluble **H)** Aβ40, **I)** Aβ42, and **J)** fibrillar amyloid. Correlation matrixes for amyloid-related proteins for **K)** prefrontal cortex and **L)** cerebellum. Statistics by ANCOVA adjusted for sex with Bonferroni’s posthoc test. Correlation matrix analyzed by Spearman correlation. *p<0.05, **p<0.01, ***p<0.001.

Clearance of Aβ peptides is mediated by apolipoprotein E (ApoE) which binds to low-density receptor 1 (LRP1). ApoE levels increased 85% in DSAD compared to AD while LRP1 was did not differ between the three groups (**Fig. 5E,F**). Aβ peptides are formed predominately as Aβ40 and the highly aggregable Aβ42. Soluble Aβ40 increased 4-fold above cognitively normal and 3-fold above AD in DSAD prefrontal cortex (**Fig. 5G**). Soluble Aβ42 had trends of increase for both AD and DSAD above cognitively normal which were not significant (**Fig. 5H**). Insoluble Aβ40 increased 3-fold above cognitively normal while insoluble Aβ42 trended upward but was not statistically significant in DSAD prefrontal cortex (**Fig. 5I,J**). Aggregation of Aβ peptides results in oligomers, proto-fibrils, fibrils, and then Aβ plaques. Fibrillar amyloid increased 2-fold in DSAD compared to cognitively normal (**Fig. 5K**)

The DSAD cerebellum had 4-fold more ADAM10 levels than cognitively normal paralleling the increase in prefrontal cortex (**Ext. Fig. 5A**). BACE1 increased with DSAD 60% from cognitively normal and 80% from AD (**Ext. Fig. 5B**). PSEN1 decreased with AD by 45% compared to cognitively normal (**Ext. Fig. 5C**). ApoE levels increased 3.5-fold in DSAD compared to cognitively normal and AD (**Ext. Fig. 5D**). LRP1 increased 45% in DSAD compared to cognitively normal and 110% higher than AD (**Ext. Fig. 5E**). Despite the increases in amyloid processing proteins in the cerebellum, no differences were observed in Aβ peptides or fibrillar amyloid (**Ext. Fig. 5F-J**). Amyloid clearance proteins ApoE and LRP1 were positively correlated in both brain regions (**Ext. Fig. 5L,M**). Soluble Aβ40 and ApoE were negatively correlated in the cerebellum suggesting increased amyloid clearance (**Ext. Fig. 5L**).

### The DSAD lipid raft has increased lipid peroxidation, reduced antioxidant enzyme defense, and increased pro-amyloidogenic processing

The lipid raft is the central signal transduction hub within all cells and is also where APP is processed. We previously described the lipid raft as a hotspot of lipid peroxidation during AD wherein antioxidant enzymes relevant to ferroptosis reside (**Fig. 6A**) [29]. Lipid raft HNE increased 2-fold in AD and 3-fold in DSAD above cognitively normal in the prefrontal cortex (**Fig. 6B**). Despite their increased lipid peroxidation DSAD rafts did not differ in lipid raft yield, cholesterol, or total protein (**Ext. Fig. 6B-D**). Lipid raft ALDH2 which clears free HNE was unaltered with AD and DSAD (**Fig. 6C**). Lipid raft GPx1, which primarily reduces hydrogen peroxide and lipid hydroperoxides cleaved from the membrane was unaltered (**Fig. 6D**). GPx1 enzyme activity was did not differ (**Fig. 6E**). GPx4 protein was decreased in the lipid raft by 60% for DSAD, more than the 30% decrease in AD (**Fig. 6F**). GPx4 activity mirrored these results with a 70% decrease in DSAD and 55% with AD (**Fig. 6G**). FSP1 also decreased with AD by 40% and trended lower with DSAD (**Fig. 6H**).

**Figure 6:**
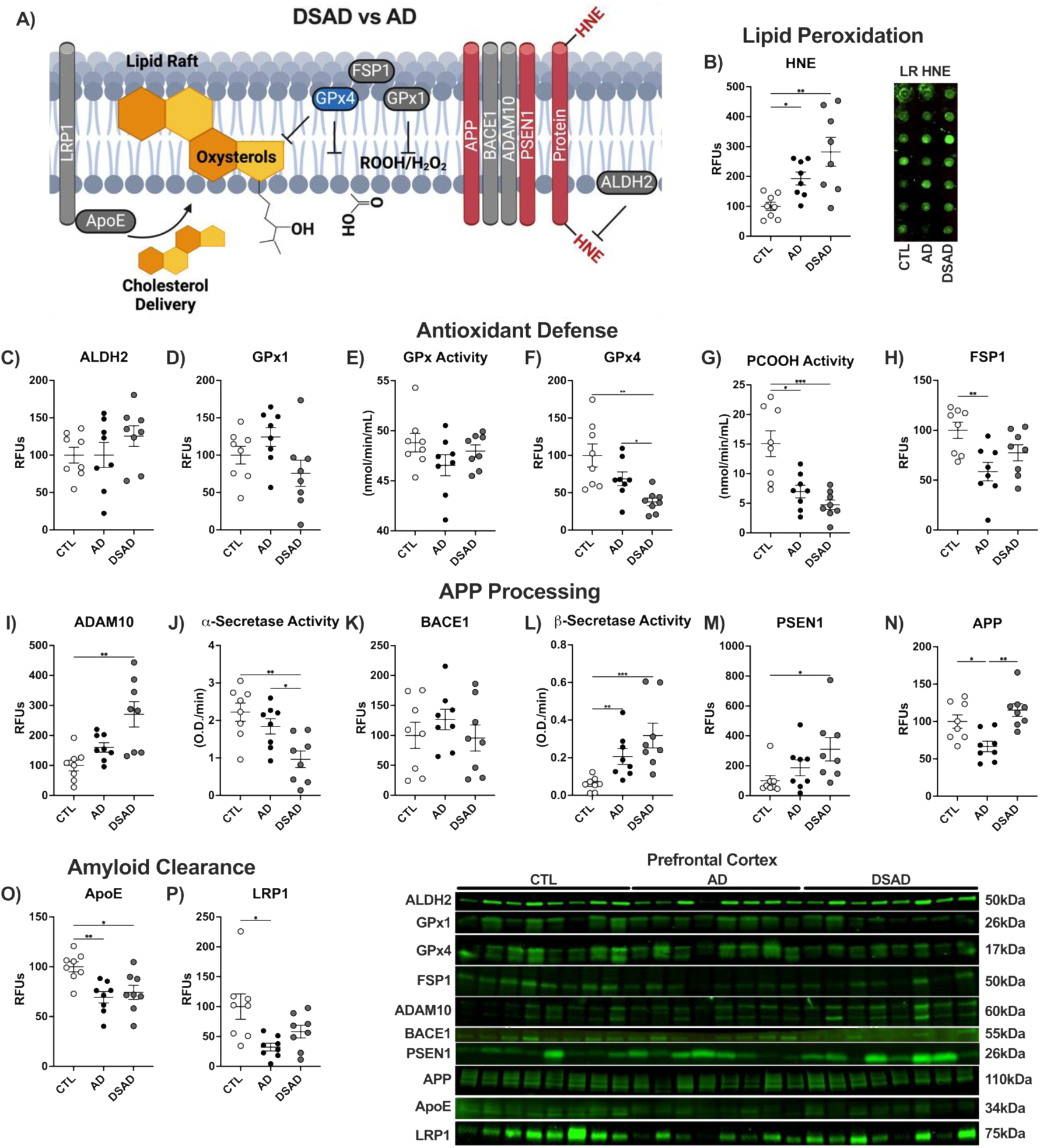
Lipid raft subcellular fraction in DSAD prefrontal cortex has increased oxidative damage and reduced antioxidant enzyme defense. **A)** Schematic representation of lipid raft changes in DSAD compared to AD. Western or dot blots shown as relative fluorescent units, or enzyme activities from lipid raft lysates for **B)** HNE, **C)** ALDH2, **D)** GPx1, **E)** GPx activity, **F)** GPx4, **G)** PCOOH activity, **H)** FSP1, **I)** ADAM10, **J)** α-Secretase activity, **K)** BACE1, **L)** β-Secretase activity, **M)** PSEN1, **N)** APP, **O)** ApoE, and **P)** LRP1. Statistics by ANCOVA adjusted for sex with Bonferroni’s posthoc test. *p<0.05, **p<0.01, ***p<0.001.

**Extended Figure 6:**
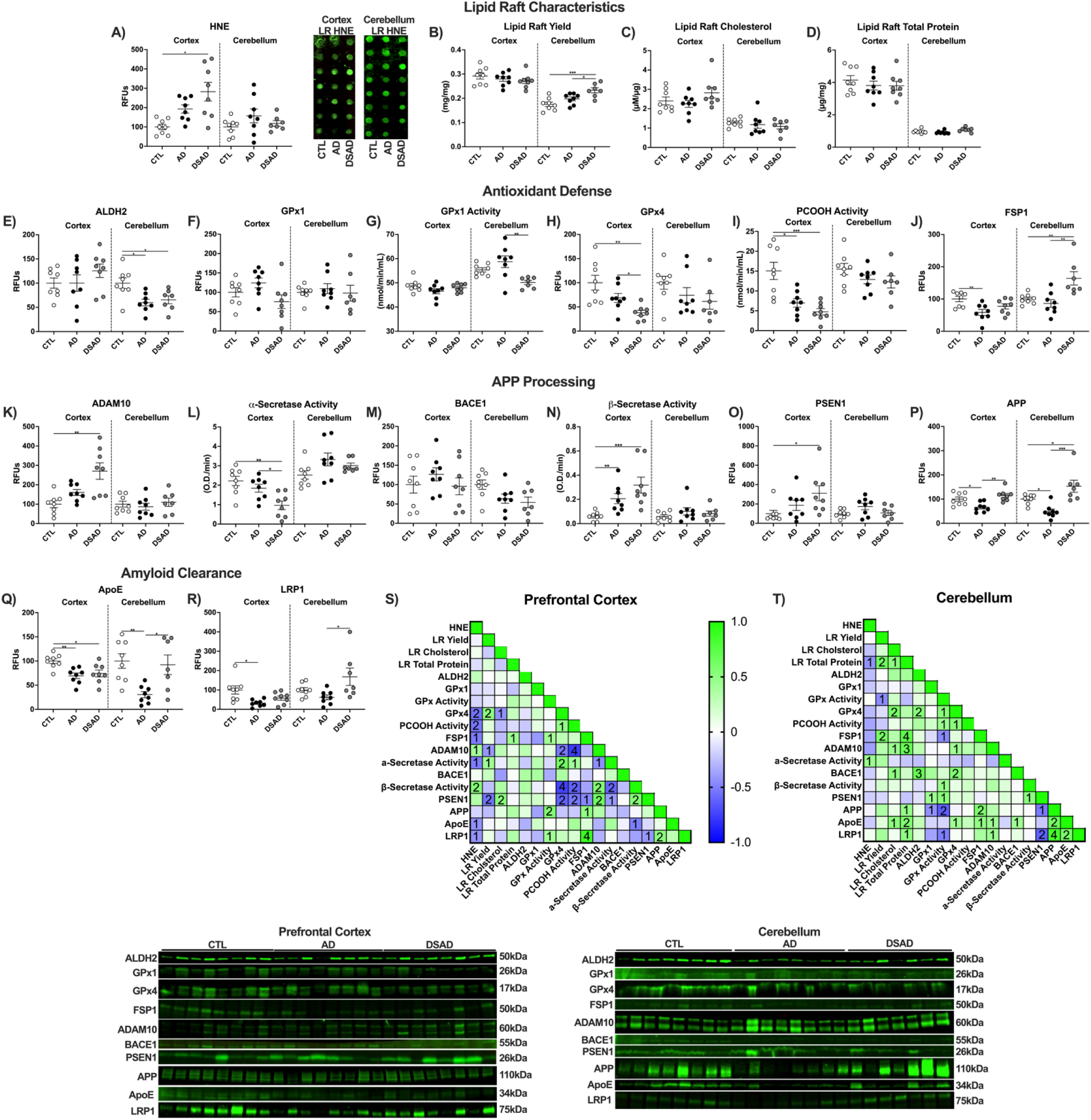
Lipid raft subcellular fraction in DSAD prefrontal cortex has increased oxidative damage and reduced antioxidant enzyme defense. Western or dot blots are shown as relative fluorescent units or enzyme activities for human prefrontal cortex or cerebellum from lipid raft lysates for **A)** HNE, **B)** yield, **C)** cholesterol, **D)** total protein, **E)** ALDH2, **F)** GPx1, **G)** GPx activity, **H)** GPx4, **I)** PCOOH activity, **J)** FSP1, **K)** ADAM10, **L)** α-Secretase activity, **M)** BACE1, **N)** β-Secretase activity, **O)** PSEN1, **P)** APP, **Q)** ApoE, and **R)** LRP1. Correlation matrix of ferroptosis-related antioxidant enzymes for **S)** prefrontal cortex and **T)** cerebellum. Statistics by ANCOVA adjusted for sex with Bonferroni’s posthoc test. *p<0.05, **p<0.01, ***p<0.001. Correlation matrix analyzed by Spearman correlation. 1p<0.05, 2p<0.01, 3p<0.001, 4p<0.0001.

Lipid rafts were examined for APP processing proteins. ADAM10 levels were 2.5-fold higher in DSAD rafts than cognitively normal (**Fig. 6I**), whereas α-secretase was 50% lower in DSAD than AD or cognitively normal (**Fig. 6J**). These opposite trends suggest that the increased ADAM10 levels may be compensatory for the low activity. BACE1 levels did not differ, but β-secretase activity increased 2.5-fold in AD and 4-fold in DSAD (**Fig. 6K,L**). Through fractionation, the localization and activity of the secretase enzymes are highest in lipid rafts rather than non-raft membranes (NRM; **Supp. Fig 1**). PSEN1 protein was 3-fold higher in DSAD than cognitively normal (**Fig. 6M**). Lipid raft APP levels changed in opposite directions with a decrease of 35% in AD and a 75% increase with DSAD above AD (**Fig. 6N**). ApoE levels decreased by 30% with AD and DSAD (**Fig. 6O**). LRP1 decreased by 75% in AD brains and trended down in DSAD compared to cognitively normal (**Fig. 6P**).

For the cerebellum, the HNE levels did not differ among the groups (**Ext. Fig. 6A**). Lipid raft yield was increased in DSAD by 35% and 20% above cognitively normal and AD respectively (**Ext. Fig. 6B**). Total cholesterol and protein did not differ between the three groups, suggesting a difference in phospholipid content with DSAD (**Ext. Fig. 6C,D**). ALDH2 was decreased by 40% with AD and 35% with DSAD below cognitively normal (**Ext. Fig. 6E**). While DSAD GPx1 protein levels did not differ, GPx1 activity decreased by 15% below AD (**Ext. Fig. 6F,G**). GPx4 protein and activity did not differ with AD and DSAD (**Ext. Fig. 6H,I**). FSP1 increased with DSAD 65% above cognitively normal and 90% above AD (**Ext. Fig. 6J**) paralleling findings in whole cell lysate (**Ext. Fig. 3F**). No difference was observed for APP processing proteins or secretase activities (**Ext. Fig. 6K-O**). Lipid raft APP decreased 50% with AD while DSAD had 2-fold more lipid raft APP than AD (**Ext. Fig. 6P**). Lipid raft ApoE decreased 70% with AD. ApoE levels were 2-fold higher in DSAD than AD (**Ext. Fig. 6Q**). LRP1 was 1.5-fold higher in DSAD compared to AD (**Ext. Fig. 6R**).

For prefrontal cortex, HNE levels were positively correlated with the pro-amyloidogenic β-Secretase activity and negatively correlated with the non-amyloidogenic α-Secretase activity. HNE and β-Secretase activity both showed negative correlations with key inhibitors of ferroptosis, GPx4, and its activity, and FSP1 in the lipid raft (**Ext. Fig. 6S**). Cerebellum did not show any of these correlations, consistent with its limited AD neuropathology (**Ext. Fig. 6T**).

### DSAD confers large transcriptomic changes from sporadic AD and cognitively normal

Transcriptomes of the prefrontal cortex and cerebellum were analyzed for changes relevant to iron metabolism and lipid peroxidation. Limited differentially expressed genes (DEGs) were found between cognitively normal and AD in prefrontal cortex. However, 1032 DEGs were identified between cognitively normal and DSAD and 939 between AD and DSAD (**S.Fig. 2A**). In the cerebellum more DEGs were found between CTL and AD with 1374, fewer than between CTL and DSAD with 278, and 2162 between DSAD and AD (**S.Fig. 2A**).

Particular genes relevant to iron are among the top 10 DEGs. The AD prefrontal cortex had increased mRNA levels for S100A4, PLXDC2, and HBB which mediate vascular remodeling and hemoglobin synthesis (**S.Fig 2B**) consistent with a relationship to MBs. Cerebellum SCARA5 was increased with AD compared to CTL and in DSAD compared to AD. SCARA5 acts as a scavenger receptor through the import of iron-loaded ferritin [42]. Upregulation of SCARA5 in AD and even more in DSAD suggests a homeostatic response for the clearance of MB-derived iron (**S.Fig. 2C**). Both brain regions had increases in the S100 family pathways consistent with the triplication of S100β with DS (**S.Fig. 2H,I**). None of the top DEGs were shared between prefrontal cortex and cerebellum.

### Does APP dosage result in increased iron?

We examined rare variants of DS with lower APP dosage for iron and hemoglobin. Mosaic Down syndrome (mT21) results from not all cells being T21 positive and partial Down syndrome (pT21) where only a segment of chromosome 21 is triplicated. Total iron in the prefrontal cortex of partial (n=1) and mosaic (n=3) T21 was 35% below DSAD (**Fig. 7A, Ext. Fig. 7A**). Tissue hemoglobin trended similarly (**Fig. 7B, Ext. Fig 7B**). Tissue iron and hemoglobin were strongly correlated (r=0.873) consistent with MB iron (**Fig. 7C**). Inclusion of AD and cognitively normal weakened this correlation as expected due to the lower tissue iron levels (r=0.632)(**Ext. Fig. 7C**).

**Figure 7:**
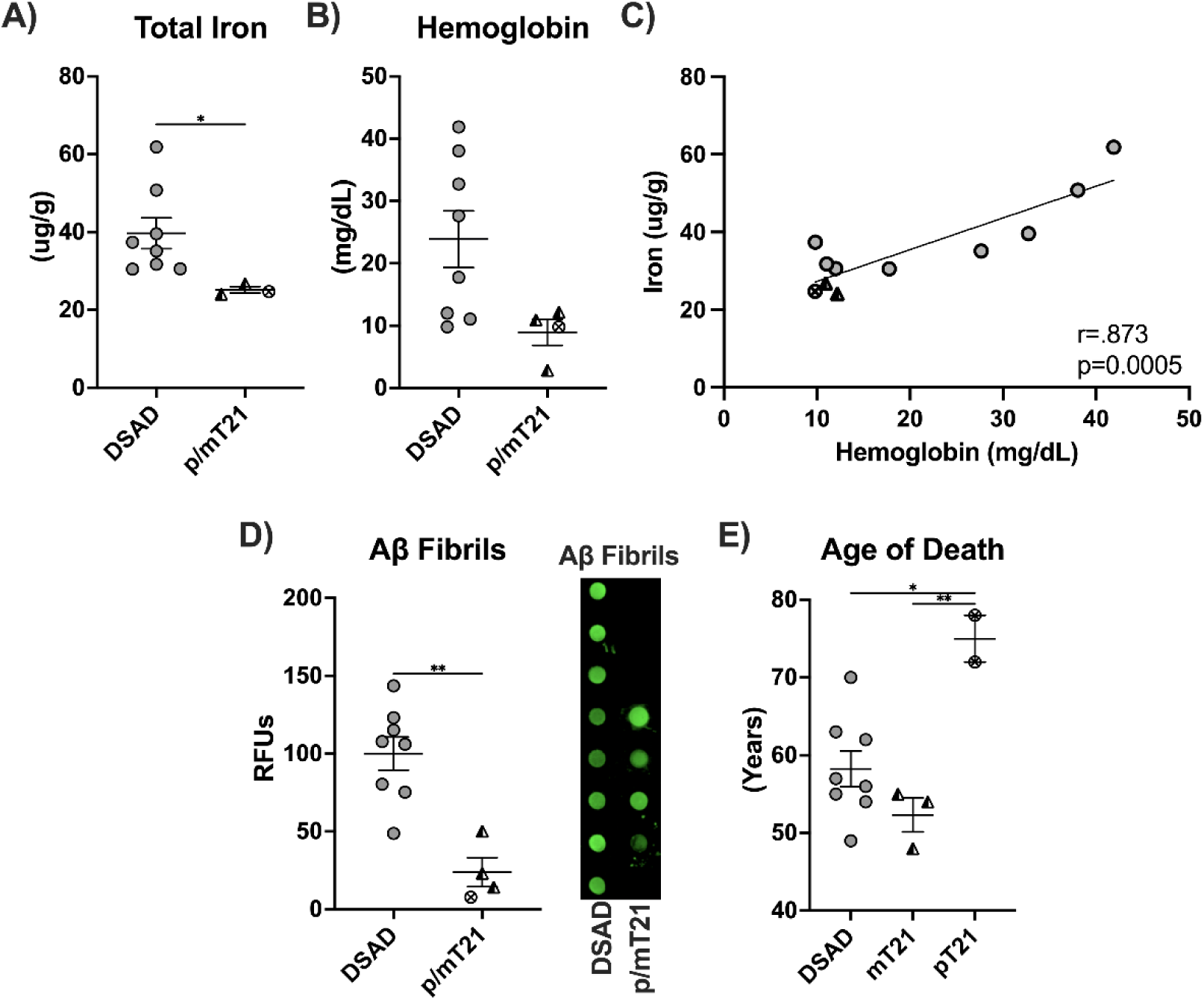
APP gene dosage and microbleed iron. Levels of **A)** total iron, **B)** tissue hemoglobin, **C)** coplot of total iron and tissue hemoglobin, dot blot shown as relative fluorescent units for **D)** Aβ fibrils, and **E)** age of death for DSAD, mosaic T21, and partial T21 cases. One additional age of death was added for pT21 without APP triplication [45]. Statistics by ANCOVA adjusted for sex with Bonferroni’s posthoc test (E) or two-tailed t-test (A, B, D) or by. Coplot by Pearson correlation. *p<0.05, **p<0.01.

**Extended Figure 7:**
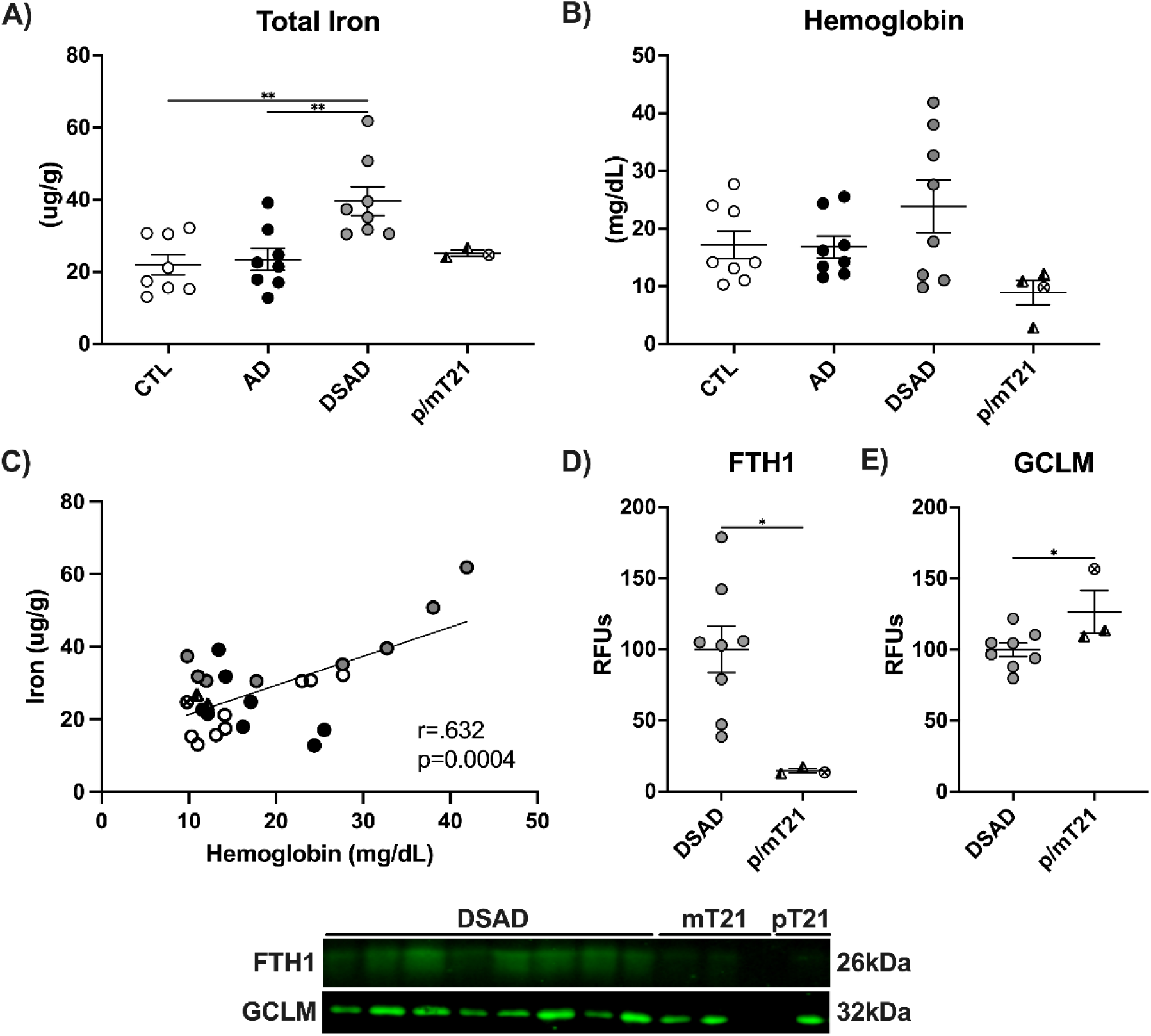
APP gene dosage and microbleed iron. Levels of **A)** total iron, **B)** tissue hemoglobin, **C)** coplot of total iron and tissue hemoglobin. Western blots are shown as relative fluorescent units for **D)** FTH1, and **E)** GCLM. Statistics by ANCOVA adjusted for sex with Bonferroni’s posthoc test (A, B) or two-tailed t-test (D, E) or by. Coplot by Pearson correlation. *p<0.05, **p<0.01.

Iron is known to seed Aβ fibrillization and Aβ fibrils are loaded with iron and other redox-active metals [43,44]. Aβ fibrils decreased by 75% in the p/mT21 prefrontal cortex compared to DSAD. This further supports the relationship between APP gene dosage and AD neuropathology (**Fig. 7D**). We next compared the age of death for DSAD, mT21 and pT21. One additional pT21 brain with documented euploidy for APP was examined [45]. pT21 conferred almost a two-decade increase in age of death compared to DSAD and mT21 (**Fig. 7E**).

Two proteins were measured in these brains due to their large differences in DSAD. Paralleling the decreases in total iron and Aβ fibrils, iron storage protein FTH1 decreased by 85% with p/mT21 (**Ext. Fig. 7D**). Finally, GCLM increased with p/mT21 by 25% (**Ext. Fig. 7E**).

## DISCUSSION

DSAD brains have more MBs than sporadic AD potentially due to the increased dosage of APP and resulting CAA. We analyzed if MB iron alters iron signaling, oxidative damage, and antioxidant enzyme defense in relation to markers of ferroptosis due to the connection between APP and iron. We previously reported that MBs precede and colocalize with amyloid plaques in FAD mice [11]. Postmortem tissues also demonstrate several key features of iron-mediated cell death referred to as ferroptosis [29]. Moreover, the lipid raft where APP is processed has extensive oxidative damage with corresponding loss of protective mechanisms during AD. Here we reexamined iron metabolism, antioxidant enzyme signaling, and lipid raft health in DSAD brains to determine if APP gene dosage worsens these outcomes. Using case and ApoE genotype-matched brain tissue, we determined that brain iron load is higher in DSAD than in AD and is associated with more lipid peroxidation and decreased antioxidant enzyme defense. The lipid raft again was a hot spot for lipid peroxidation with levels above sporadic AD and lesser antioxidant enzyme defense for DSAD. Comparisons with rare DS variants of partial or mosaic trisomy confirmed that brain iron levels increase with triplicated APP.

Amyloid neuropathology increased in DSAD as expected, matching the reported Braak staging and earlier reported deposition of amyloid plaques in DS brains. However, the lipid raft revealed increased ADAM10 levels which primarily process APP through the non-amyloidogenic pathways. This may be due to the increased APP levels or because of the enhanced oxidative damage to DSAD lipid rafts. Electrophilic adduct modification of proteins occurs by the lipid peroxidation product HNE on amino acids [46], or nitrotyrosine on tyrosine residues; tyrosine itself does not conjugate with HNE [47]. Amino acids that span membranes, or are near membrane surfaces, are most vulnerable to conjugation by carbonyls. HNE adducts increase BACE1 activity, while NT may diminish enzyme activity [30,48]. We see opposing changes for alpha and beta-secretase activity in the DSAD raft, potentially from oxidative modification. ADAM10 contains the largest number of vulnerable residues which may explain the decrease in enzymatic activity despite increased protein levels. The hypothesis of increased ADAM10 oxidation is further supported by the lowered antioxidant enzyme defense and increased lipid peroxidation in the lipid raft.

Many enzymes that mediate the repair of oxidized membrane lipids are GSH-dependent. GCLM is required for GSH formation in mammals and is needed for the rate-limiting step in GSH formation. While GSH was not measured due to its rapid degradation in postmortem samples, the decrease in GCLM in AD and DSAD is suggestive of reduced brain GSH which was confirmed elsewhere by MRI [49,50]. Furthermore, plasma GSH is a third lower in DS children than non-DS [51] which enhances oxidative damage during preadolescence. We found increased GPx4 and FSP1 in whole-cell lysates; these membrane enzymes are key players in preventing ferroptosis [52]. However, lipid rafts had decreased levels of both GPx4 and FSP1 in AD that were even larger in DSAD for GPx4. The levels of GPx4 protein, which reduces phospholipid hydroperoxides, matched the enzyme activity levels of PCOOH reduction. Both GPx4 and FSP1 require localization signals to direct them to membranes which may explain the reduced presence of these proteins in the lipid raft during AD [29,53,54]. The increase in GPx4 and FSP1 protein is potentially compensatory, due to their lowered localization to lipid rafts in an attempt to force them to localize or to reduce oxidized phospholipids peripheral to the lipid raft.

The connection between AD and DS has largely been attributed to APP on chromosome 21 because of the increase in Aβ neuropathology by age 30 [55,56]. However, other triplicated genes such as BACH1, a competitive inhibitor of antioxidant enzyme gene transcription, may play a large role in propagating lipid peroxidation in the prodromal phases of DSAD. BACH1 is degraded in the presence of heme [57], allowing Nrf2 to bind to the HMOX1 and other promotors of antioxidant enzyme genes for heme degradation, iron storage, and antioxidant enzyme defense. A decrease in BACH1 was observed despite its triplication, explained by the increase in total iron and HMOX1 in the prefrontal cortex. Moreover, we see increased iron storage and import with a simultaneous drop in heme import, suggestive of excess heme. This increase in heme iron may be MB derived. These differences are absent from the cerebellum which has less AD neuropathology and fewer MBs than the cortical and subcortical regions [58]. Collectively these data implicate the ferroptotic changes in DSAD attributed to MB iron in AD vulnerable brain regions.

Another DSAD study showed BACH1 was increased, which may be attributed to Brodmann area differences [59]. This study also found BACH1 increased in DS patients compared to young controls which would stifle Nrf2-mediated gene induction. This would result in difficulties maintaining homeostatic levels of iron and repair of lipid peroxidation accelerating pathological events. Regardless, the pT21 case used within the manuscript has deletion of APP but not BACH1 on the triplicated chromosome [60] which was enough to attenuate cortical iron levels and increase GCLM with a later age of death of 72 years. The ‘Prasher case’ had a deletion of both APP and BACH1 and a slightly later age of death at 78 years. Both of these brains had low Braak staging and ‘normal’ or no detectable amyloid levels. While these two rare DS patients alone are not sufficient for strong conclusions, they suggest APP gene dosage may be linked to iron levels and overall lifespan aside from the clear implications in amyloid neuropathology. Contrarily, the mT21 brains have lower APP gene dosage due to not all cells being T21 positive, but the same average age of death as DSAD despite lower iron, tissue hemoglobin, and fibrillar amyloid levels highlighting unknown complexities that extend beyond APP gene dosage alone.

The current study is limited by the sample size which is underpowered to separately consider sex and differences by ApoE allele. While the focus was to examine differences between sporadic and DSAD, we had an insufficient number of DS brains without AD to include for comparison. This makes it unclear which changes are due to DS-related or AD neuropathology. For these reasons, emphasis was placed on differences observed in both AD and DSAD. DSAD patients have received little support through clinical trials for the treatment of AD. One trial is currently ongoing using Donanemab which targets only pyroglutamate amyloid, which constitutes a smaller fraction of the total amyloid species [61]. Despite only targeting roughly 10-25% of the amyloid pool Donanemab has promise likely due to this selectivity. Future studies on MBs in the various types of AD must consider the link between brain iron and amyloid. We recently showed that deferoxamine, an iron chelator, reduces brain iron in wildtype mice and Aβ fibrils in FAD mice strengthening the connection between iron and amyloid pathology [29,62]. Does more aggressive removal of iron-laden amyloid plaques lead to massive oxidative damage from liberated iron?

---

**Supplemental Table 1:** Patient information for the tissue used in this study.

| Diagnosis | Genotype | Braak | Age | Sex | PMI | ADRC |
| --- | --- | --- | --- | --- | --- | --- |
| CTL | ApoE3,3 | 0 | 82 | Male | 9 | USC |
| CTL | ApoE3,3 | 0 | 93 | Male | 12 | USC |
| CTL | ApoE3,3 | 0 | 93 | Male | 3.75 | USC |
| CTL | ApoE3,3 | 0 | 76 | Male | 11.25 | USC |
| CTL | ApoE3,3 | 0 | 99 | Female | 9 | USC |
| CTL | ApoE3,3 | 0 | 91 | Female | 8.75 | USC |
| CTL | ApoE3,3 | 1 | 95 | Female | 3.25 | USC |
| CTL | ApoE3,3 | 0 | 85 | Female | 7 | USC |
| AD | ApoE3,3 | 4 | 87 | Male | 4.75 | USC |
| AD | ApoE3,3 | 5 | 88 | Male | 6.75 | USC |
| AD | ApoE3,3 | 3 | 97 | Male | 5.25 | USC |
| AD | ApoE3,3 | 5 | 76 | Male | 9.75 | USC |
| AD | ApoE3,3 | 5 | 81 | Female | 7.5 | USC |
| AD | ApoE3,3 | 6 | 66 | Female | 17 | USC |
| AD | ApoE3,3 | 5 | 89 | Female | 1.5 | USC |
| AD | ApoE3,3 | 4 | 94 | Female | 10.5 | USC |
| DSAD | ApoE3,3 | 6 | 54 | Male | 4.3 | UCI |
| DSAD | ApoE3,3 | 6 | 49 | Male | 4 | UCI |
| DSAD | ApoE3,3 | 6 | 55 | Male | 3.18 | UCI |
| DSAD | ApoE3,3 | 6 | 70 | Male | 3.87 | UCI |
| DSAD | ApoE3,3 | 6 | 57 | Female | 4.25 | UCI |
| DSAD | ApoE3,3 | 6 | 63 | Female | 3.58 | UCI |
| DSAD | ApoE3,3 | 6 | 62 | Female | 3.8 | UCI |
| DSAD | ApoE3,3 | 6 | 56 | Female | 2.92 | UCI |
| mDS | ApoE3,3 | 6 | 54 | Male | 4.5 | UCI |
| mDS | ApoE3,3 | 3 | 48 | Female | 18.35 | UCI |
| mDS | N/A | 1 | 55 | Male | N/A | UCI |
| pDS | ApoE2,3 | 3 | 72 | Male | 4.87 | UCI |

**Supplemental Figure 1:**
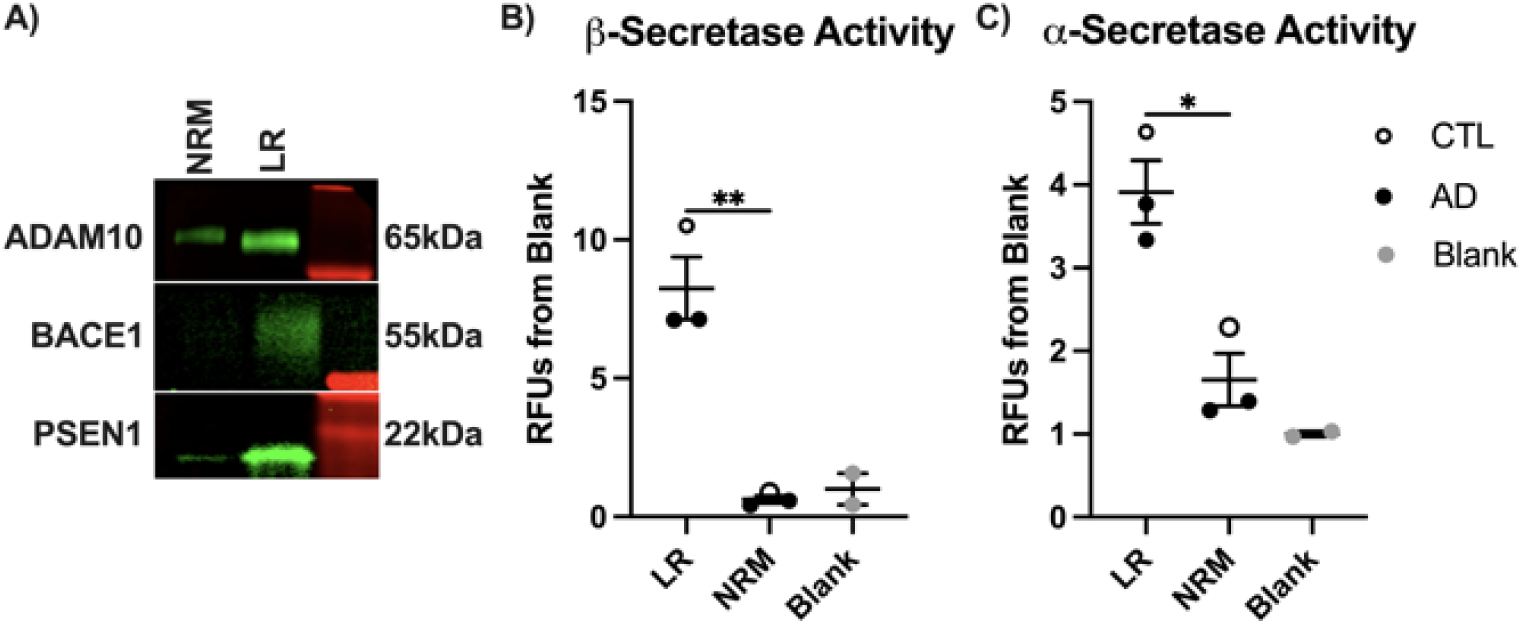
Validations of Secretase localization and activities. **A)** comparison of NRM and LR fractions for ADAM10, BACE1, and PSEN1 by Western Blot in human prefrontal cortex. Comparison of NRM, LR, and blank wells (all reagents except sample) for **B)** β-secretase, and **C)** α-secretase activities presented as relative fluorescence from blank in human prefrontal cortex. t-test, *p<0.05, **p<0.01.

**Supplemental Figure 2:**
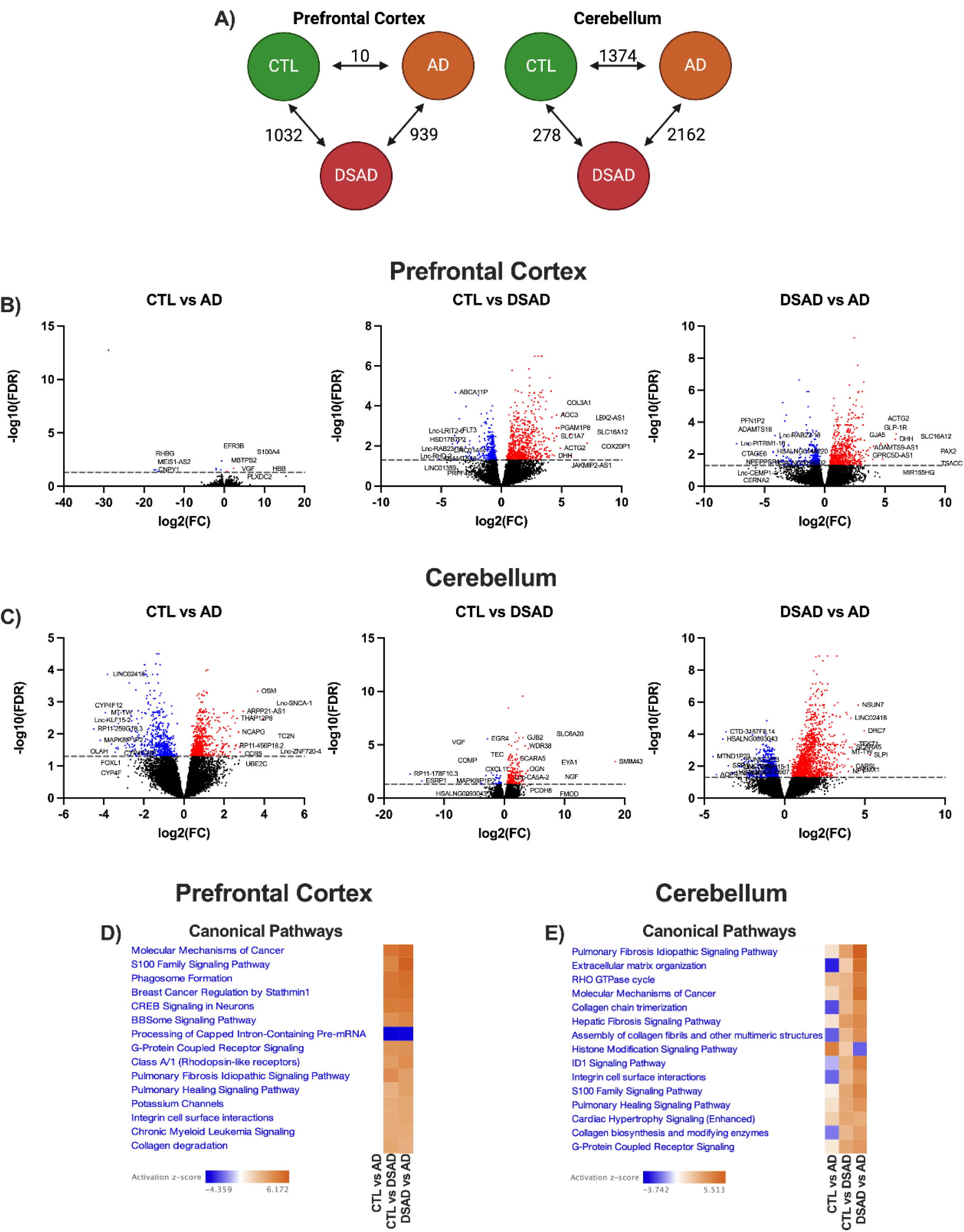
Transcriptional changes with sporadic AD and DSAD. **A)** Shared differentially expressed genes (DEGs) between cognitively normal, AD, and DSAD for prefrontal cortex and cerebellum. Volcano plots showing the top 10 up and down-regulated genes for CTL vs AD, CTL vs DSAD, and DSAD vs AD for **B)** prefrontal cortex and **C)** cerebellum. The top 15 identified pathways altered in **D)** prefrontal cortex, and **E)** cerebellum.

## Consent Statement

All human subjects provided informed consent.

## Acknowledgments

The authors are grateful to Liying Zhao for performing ICP-MS measurements, and Carol Church and Kymry Jones for assistance with postmortem tissues. The authors appreciate the helpful discussions of brain samples with Carol Miller (USC) and C. Dirk Keene (UW). Tissue for this study was obtained from the USC ADRC Neuropathology Core, NIA-AG066530. The UCI-ADRC is funded by NIH/NIA Grant P30-AG066519. J. Silva and E. Head were supported by P30-AG066519. Lab studies were supported by NIH grants to CEF (R01-AG051521, P50-AG05142, P01-AG055367) and Cure Alzheimer’s Fund; Simons Collaboration on Plasticity in the Aging Brain grant SF811217 to BB; R01AG079806 from the NIA, Larry L. Hillblom Foundation Grant 2022-A-010-SUP, the Glenn Foundation for Medical Research and AFAR Grant for Junior Faculty Award, and Navigage Foundation Award to RHS; T32AG052374 and R01AG079806-02S1 to GG; and T32-AG000037 (PI: Eileen Crimmins) to MAT. Brain specimens were obtained from ADRC Tissue Cores: USC (P50-AG005142, AG066530); University of California Irvine (P30-AG066519); UW (P30-AG066509; U01-AG006781). Venn diagrams were made with Venny 2.1.0 and schematics were made using BioRender.

## Ethics Declarations

All of the authors declare no competing interests.

## Data Availability

The authors declare that the data supporting the findings of this study are available within the paper and its Supplementary Information files. Should any raw data files be needed in another format they are available from the corresponding author upon reasonable request.

